# Variant-specific changes in persistent or resurgent Na^+^ current in *SCN8A*-EIEE13 iPSC-derived neurons

**DOI:** 10.1101/2020.01.16.909192

**Authors:** Andrew M. Tidball, Luis F. Lopez-Santiago, Yukun Yuan, Trevor W. Glenn, Joshua L. Margolis, J. Clayton Walker, Emma G. Kilbane, Lori L. Isom, Jack M Parent

**Author notes:** Co-corresponding authors, Corresponding authors: Jack M Parent, Lori L. Isom.

## Abstract

Missense variants in the voltage-gated sodium channel (VGSC) gene, *SCN8A*, are linked to early-infantile epileptic encephalopathy type 13 (EIEE13). EIEE13 patients exhibit a wide spectrum of intractable seizure types, severe developmental delay, movement disorders, and elevated risk of sudden unexpected death in epilepsy (SUDEP). The mechanisms by which *SCN8A* variants lead to epilepsy are poorly understood, although heterologous expression systems and mouse models have demonstrated altered sodium current (I_Na_) properties. To investigate these mechanisms using a patient-specific model system, we generated induced pluripotent stem cells (iPSCs) from three patients with missense variants in *SCN8A*: p.R1872>L (P1); p.V1592>L (P2); and p.N1759>S (P3). Using small molecule differentiation into excitatory neurons, iPSC-derived neurons from all three patients displayed altered I_Na_. P1 and P2 had elevated persistent I_Na_, while P3 had increased resurgent I_Na_ compared to controls. Further analyses focused on one of the patients with increased persistent I_Na_ (P1) and the patient with increased resurgent I_Na_ (P3). Excitatory cortical neurons from both patients had prolonged action potential (AP) repolarization and shorter axon initial segment lengths compared to controls, the latter analyzed by immunostaining for ankyrin-G. Using doxycycline-inducible expression of the neuronal transcription factors Neurogenin 1 and 2 to synchronize differentiation of induced excitatory cortical-like neurons (iNeurons), we investigated network activity and response to pharmacotherapies. Both patient neurons and iNeurons displayed similar abnormalities in AP repolarization. Patient iNeurons showed increased burstiness that was sensitive to phenytoin, currently a standard treatment for EIEE patients, or riluzole, an FDA-approved drug used in amyotrophic lateral sclerosis and known to block persistent and resurgent I_Na_, at pharmacologically relevant concentrations. Patch-clamp recordings showed that riluzole suppressed spontaneous firing and increased the AP firing threshold of patient-derived neurons to more depolarized potentials. Our results indicate that patient-specific neurons are useful for modeling EIEE13 and demonstrate *SCN8A* variant-specific mechanisms. Moreover, these findings suggest that patient-specific iPSC neuronal disease modeling offers a useful platform for discovering precision epilepsy therapies.

## Introduction

During the recent explosion in exome sequencing of epilepsy patients, *SCN8A* was linked to early infantile epileptic encephalopathy type 13 (EIEE13) (Veeramah *et al.*, 2012; Epi4K and Investigators, 2013; Ohba *et al.*, 2014; Larsen *et al.*, 2015). EIEE13 patients have a wide spectrum of seizure types with seizure onset occurring on average at 5 months of age (Larsen *et al.*, 2015). Developmental delays, lack of speech, and hypotonia are nearly universal. Seizures are typically severe in EIEE13 patients, refractory to antiseizure medications (ASMs), and lead to a high risk of sudden unexpected death in epilepsy (SUDEP), findings that have been recapitulated in mouse models (Veeramah *et al.*, 2012; Wagnon *et al.*, 2014; Kong *et al.*, 2015; Larsen *et al.*, 2015; Bunton-Stasyshyn *et al.*, 2019).

*SCN8A* encodes the voltage-gated sodium channel (VGSC) α subunit, Na_v_1.6, that is the most abundant VGSC in the central nervous system, where it is primarily localized to axon initial segments (AIS) and nodes of Ranvier (Gasser *et al.*, 2012). Nearly all *SCN8A* variants linked to EIEE13 thus far are *de novo* missense mutations resulting in single amino acid substitutions (Wagnon and Meisler, 2015), although one patient inherited the mutation from a mosaic parent (Larsen *et al.*, 2015). Data from a knock-in mouse model and from heterologous expression of several mutations in cultured cells implicate a variety of potential sodium current (I_Na_) abnormalities in disease pathogenesis, including slowed inactivation kinetics (Wagnon *et al.*, 2016), increased persistent I_Na_ (Veeramah *et al.*, 2012; Wagnon *et al.*, 2016; Lopez-Santiago *et al.*, 2017; Ottolini *et al.*, 2017), and increased resurgent I_Na_ (Patel *et al.*, 2016; Ottolini *et al.*, 2017). The predicted outcomes of these changes in I_Na_ properties are delayed action potential (AP) repolarization and ectopic firing. Consistent with this, recordings from the *Scn8a^N1768D/+^* EIEE13 mouse model showed early after depolarization (EAD)-like AP waveforms in CA1 hippocampal neurons (Lopez-Santiago *et al.*, 2017). A complete understanding of *SCN8A* mutation-specific mechanisms would necessitate the production of multiple mouse models. Moreover, because data from mice often do not translate to humans, it is desirable to test ASM efficacy in other models such as cultured human cortical neurons, which is now feasible in patient-derived cells using induced pluripotent stem cell (iPSC) technology.

While mouse and heterologous systems are useful, we sought to perform mechanistic studies in a human neuronal system that allows rapid testing of potential EIEE13 mutation-specific effects. To this end, we generated iPSC lines from 3 EIEE13 patients. Cortical excitatory patient neurons differentiated from the iPSCs exhibited elevated persistent or resurgent I_Na_ compared to control neurons, similar to findings in mouse and heterologous cell culture models. EIEE13 patient neurons also showed prolonged AP repolarization and shorter AIS lengths. To reduce experimental variability inherent in small molecule cortical neuron differentiations, we generated doxycycline-inducible neurons (iNeurons) from the same iPSC lines. iNeurons displayed increased network bursting, as assessed by burst duration and percentage of spikes in bursts. Genotypic differences in network activity were rescued by administration of phenytoin or riluzole, the latter a drug with relative selectivity for persistent and resurgent I_Na_.

## Materials and methods

### Induced pluripotent stem cell reprogramming

Skin punch biopsies were obtained from 3 EIEE13 patients (P1-3) and two healthy controls (C1 and C4) without known genetic disorders with consent under a protocol approved by the Institutional Review Board of Michigan Medicine and reprogrammed by previously published methods (Tidball *et al.*, 2017). Two additional control lines used in this study were previously reported as CC1 (now C2) and *CHD2* WT/WT2 (now C3) from (Tidball *et al.*, 2016; Tidball *et al.*, 2017), respectively, and were reprogrammed using identical methods. See **Supplemental materials** for further details of reprogramming and cell line validation.

### CRISPR genome-editing with simultaneous reprogramming

We have previously published methods combining CRISPR genome-editing and iPSC reprogramming (Tidball *et al.*, 2017; Tidball *et al.*, 2018). See **Supplemental materials** for further details describing the generation of the P2 rescue iPSC line.

### Excitatory neuron differentiation

We utilized a modification of the Shi *et al.* technique for differentiating iPSCs into excitatory cortical neurons. The neuronal cultures were transduced with lentivirus containing a mouse *Camk2a* promoter-driven GFP to identify mature neurons (Shcheglovitov *et al.*, 2013; Nehme *et al.*, 2018). See **Supplemental material** for further details.

### Generation and differentiation of stable doxycycline (dox)-inducible *Ngn1/2*(iNeuron) iPSC lines

Established iPSC lines were transfected using Mirus LT-1 with 2 TALEN plasmids (pZT-C13-R1 and pZT-C13-L1, gifts of Jizhong Zou; addgene.org: #52638 and 52637, respectively) targeting the safe-harbor-like locus, *CLYBL,* and a targeting plasmid containing the dox-inducible promoter system which controls *Neurogenin1* (*Ngn1*) and *Ngn2* expression, and constitutively active *mCherry* (pUCM-CLYBL-*Ngn1/2*, a gift of Dr. Michael Ward). Pure mCherry+ clonal lines were obtained by manual selection. To differentiate into iNeurons, cells were grown for 72 hours in mTeSR1 media containing doxycycline. Cells were then replated onto PEI/laminin coated dishes in 3N medium containing doxycycline. On day 8, the medium was replaced with BrainPhys medium containing N2 and SM1 supplements as well as BDNF and GDNF. See **Supplemental material** for further details.

### Immunostaining

Neurons grown on Nunc™ Lab-Tek™ III 8-well chamber slides (Thermo Scientific) were fixed in paraformaldehyde for 30 min at room temperature (RT). After permeabilization with 0.2% Triton-X 100 for 20 min at RT, cells were incubated in PBS containing 5% normal goat serum with 1% BSA and 0.05% Triton-X100 for 1 h at RT. Cell were incubated in primary antibody overnight (see **Supplemental Table 2** for antibodies and dilutions) in the same blocking buffer at 4 ºC, washed 4 times in PBS with 0.05% Tween-20 (PBST), and incubated for 90 min with secondary antibody. Cells were washed 3 times in PBST and incubated with bisbenzimide for 5 min. After additional PBS washes, coverslips were mounted on slides with Glycergel mounting medium (Agilent Dako). Images were obtained on a Leica SP5 upright DMI 6000 confocal microscope.

### AIS imaging and measurement

Neurons were differentiated and immunostained on day 21 as described above with antibodies for ankyrin-G and microtubule associated protein 2ab-isoforms (MAP2ab). AIS were imaged on a Leica SP5 upright DMI 6000 confocal microscope using a water-immersion 63x objective. Images were taken at 1 Airy unit for the pinhole size with overlapped Z-series steps covering the entire AIS thickness. Max projection images were obtained and exported as TIFF files, then imported into ImageJ. Segmented lines were overlaid by a blinded observer. The plot profiles (intensity of each pixel on the line) were exported and uploaded to a MATLAB program designed for determining AIS length (Dr. Eugene Katrukha at Utrecht University). In short, this program performs a smoothening function, normalizes the intensity data, and applies a threshold of 40% to determine the ends of the AIS (Yau *et al.*, 2014).

### Electrophysiological recordings

Single GFP+ (labeled with a lentiviral *Camk2a*-GFP reporter) iPSC-derived neurons were analyzed by whole-cell patch clamp recordings. Voltage clamp recordings were performed in the standard whole-cell configuration, using previously described conditions (Liu *et al.*, 2013). All experiments were carried out at RT (21-22 °C). See **Supplemental material** for further details.

### Multielectrode array (MEA) recordings

On day 3 of dox treatment, differentiating iNeurons were plated onto PEI/laminin coated Axion 96-well MEA recording plates at 1.5 × 10^5^ cells in a 5 μL drop on the electrode grid (8 electrodes/well) of each well. Cells were grown as described above with half-volume media changes 3 times/week. Every 2-4 days a 5-min recording was performed at 37 °C after a 5-min equilibration time on the MEA. At 30 days after dox treatment, the MEA plates were used for ASM testing. Due to differences in the overall activity of individual plates, the definition of a network burst was changed from default settings to reduce false positive and false negative network burst detections (Matsuda *et al.*, 2018). Therefore, we employed an iterative process of altering the minimum percentage of electrodes participating (from 25-50%, or 2-4 electrodes) and the minimum number of spikes (from 10-100 spikes) for each plate on the most active recording day (typically day 30). A maximum interspike interval (ISI) of 75 ms was used to define bursts for all analyses.

Between days 30-33, drug testing was performed on MEA plates that did not show signs of peeling or excessive cell death (shown by loss of overall activity or lack of network bursting in all wells). Concentrated (10x) ASMs were added to the plate in a 10% volume (20 μL) with 1% DMSO, or DMSO alone (control). Thus, final concentrations were 1x of the expected concentration with 0.1% DMSO. Plates were allowed to equilibrate for 5 min on the Maestro MEA followed by a 10-min baseline measurement. The plate was immediately removed, and the drugs were added. Each drug/concentration was added to 1 row yielding 6 wells used per line. The plate was placed back on the MEA and allowed to equilibrate for 5 min followed by a 10-min experimental recording. When performing drug-testing analyses, the percentage of spikes in network bursts values were only used from wells that had a pre-treatment level of at least 5% spikes in network bursts. For all other measures, wells with<0.01 Hz mean firing rate (MFR) were removed from analyses. Each well of the drug-testing MFR data was normalized to the pretreatment level to account for large plate-to-plate variability in overall activity.

### Statistical analyses

Results are expressed as mean ± SEM except for the iNeuron AP dynamics, which are displayed as 95% confidence intervals (CI). Data were analyzed and organized using Excel (Microsoft). Prism (GraphPad) was used for generating graphs and statistical analyses. Sodium current, AIS length, and AP dynamic data were analyzed by the non-parametric Kruskal-Wallis test with uncorrected Dunn’s post-test. For MEA data analyses, when no data points were missing, we used repeated measures two-way ANOVA. Post-hoc analyses were performed by two-stage linear step-up procedure for multiple comparison controlling for false discovery rate (Benjamini *et al.*, 2006).

## Results

### EIEE13 patient-derived neurons show variant-specific differences in I_Na_ properties

To understand the disease mechanisms of individual EIEE13 patient *SCN8A* variants in a human neuronal model, we generated integration-free iPSCs by episomal transfection of fibroblasts from 3 EIEE13 patients (**Fig. 1A; Table 1**) and 2 non-epileptic controls, C1 and C4 (**Table 1**). Patient variants were confirmed in each iPSC line by PCR of genomic DNA followed by Sanger sequencing (**Supplemental Fig. 1A**). We used two additional control lines, C2 and C3, (**Table 1**) that have been previously published (Tidball *et al.*, 2016; Tidball *et al.*, 2017). Each line used for subsequent experiments was negative for episomal reprogramming vector integration, had no observable genomic abnormalities as tested by either g-band karyotype or SNP chip microarray analyses, and expressed markers of pluripotency (**Supplemental Fig. 1B, C and data not shown**). A subset of the iPSC lines (2 patients and 1 control) were differentiated to embryoid bodies, resulting in the formation of 3 germ layers as assessed by immunofluorescence microscopy (**Supplemental Fig. 1D**).

**Table 1.** Patient information. Information for patients and controls from which the iPSC lines were derived are presented including, age at biopsy, sex, *SCN8A de novo* variant, seizure type, medications, and other findings.

| Table 1. information for patients and controls from which the iPSC lines were generated. |  |  |  |  |  |  |  |
| --- | --- | --- | --- | --- | --- | --- | --- |
|  | Seizure onset | Age at biopsy/sex | Mutations | Seizure type | Other findings | Previous antiseizure medications | Current medications |
| **Patient 1 (P1)** | 1 mo | 16 yo F | p.R1872>L | myoclonic at first, now gelastic | intellectual disability, pancerebellar atrophy and ataxia | Keto, PB, CZP, VPA, LTG, PRM, GAB, OXC, LEV, ZON, FBM | PHT, CLB, TPM, vagal nerve stimulator |
| **Patient 2 (P2)** | 5 mo | 9 mo F | p.V1592>L | Focal onset with eye deviation | developmental delays, hypotonia, apnea |  | OXC, infrequent seizures |
| **Patient 3 (P3)** | 10 mo | 7 yo F | p.N1759>S | myoclonic jerks, altered mental status | encephalopathy, developmental delays, ataxia | LAC, PHT, OXC | LEV |
| **Patient 4 (P4)\*** | neonatal | 1 yo M | p.V1757>I | tonic | hyperekplexia, bradycardia, apnea | ZON, LOR | PHT, OXC, GBP, CZP |
| **Control 1 (C1)** |  | 36 yo F |  |  |  |  |  |
| **Control 2 (C2)** |  | 17 yo F |  |  |  |  |  |
| **Control 3 (C3)** |  | Newborn M |  |  |  |  |  |
| **Control 4 (C4)** | Mother of Patient-1 | 47 yo F |  |  |  |  |  |
| * Neuronal data for this patient is not presented in this publication |  |  |  |  |  |  |  |
| Keto: ketogenic diet, PB: phenobarbital, CZP: clonazepam, VPA: valproic Acid, LTG: lamotragine, PRM: primidone, GAB: gabazine, OXC: oxcarbazepine, LEV: levetiracetam, ZON: zonisamide, FBM: felbamate, PHT: phenytoin, CLB: clobazam, TPM: topiramate, LOR: lorazepam, RFM: rufinamide. GBP: gabapentin |  |  |  |  |  |  |  |

**Figure 1.**
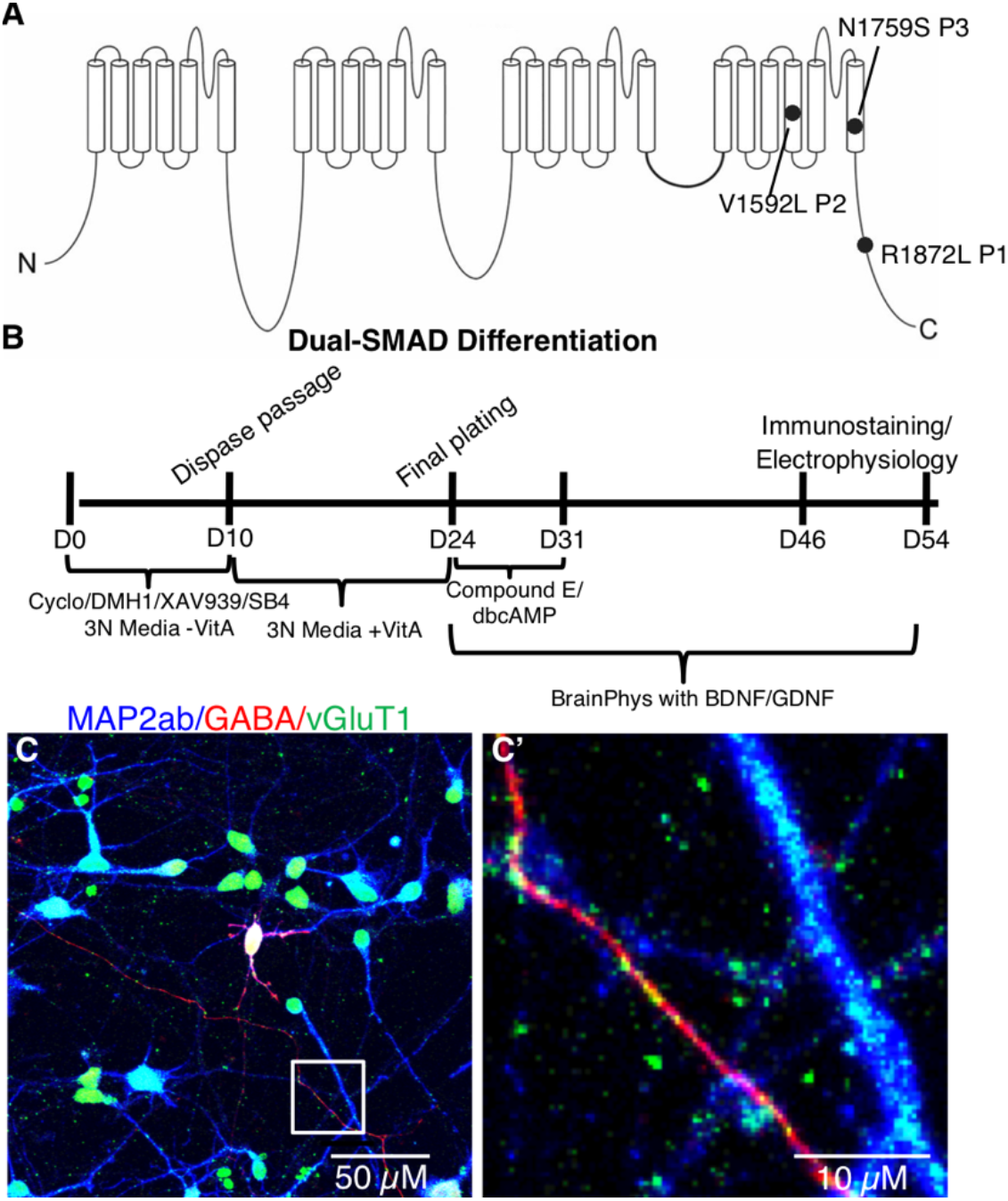
Patient variants and glutamatergic neuronal differentiation. (**A**) Diagram of the *SCN8A* gene with *de novo* missense variants from the three patients included in this study. (**B**) Differentiation protocol for generating excitatory cortical neurons from iPSCs using dual-SMAD inhibition combined with WNT and SHH inhibition. (**C**) Mature iPSC-derived neuronal cultures labeled with MAP2ab (blue), GABA (red), and vGluT1 (green). (**C’**) Enlarged image of a section of panel **C** highlighting vGLUT1-labeled (green) puncta apposed to GABA- (red) and MAP2ab-positive (blue) neuronal processes.

We differentiated the iPSCs into cortical excitatory neurons using a dual-SMAD inhibition protocol for specifying PAX6^+^ neuroepithelium (**Fig. 1B**). Our use of DMH1 for brain morphogenic protein (BMP) inhibition instead of dorsomorphin or Noggin was based on its more specific inhibition of activin receptor-like kinase-2 (ALK2) (Neely *et al.*, 2012). We also added the Wnt-antagonist XAV939, which is known to increase the forebrain identity of dual-SMAD differentiated neurons (Maroof *et al.*, 2013), as well as cyclopamine to inhibit any endogenous sonic hedgehog (SHH) signaling and, thereby, decrease the number of subpallial GABAergic neurons generated. Using immunocytochemistry, nearly all patient and control neurons were positive for vesicular glutamate transporter 1 (vGLUT1) with ~10% GABAergic neurons, which often double-labeled with both vGLUT1 and GABA (**Fig. 1C, C’**), consistent with recent studies that reported several cell types in the rodent brain that co-express these two markers (Perreault *et al.*, 2012; Root *et al.*, 2018). About 30% of cells expressed CTIP2 at day 43 of differentiation (data not shown), as was reported in the original publication of the protocol we adapted (Shi *et al.*, 2012). We also performed qRT-PCR for *SCN8A, SCN1A, SCN1B,* and *SCN4B* and found no differences between the patient and control groups (**Supplemental Fig. 2**). The mRNA expression for *SCN4B* was not detectable in our cultures.

We next measured I_Na_ properties of EIEE13 patient and control iPSC-derived neurons. Whole-cell voltage clamp recordings were performed on isolated neurons labeled with GFP driven by the *Camk2a* promoter, a marker of excitatory cortical neurons that has been shown to identify neurons with more mature electrophysiological characteristics (**Fig. 2F**) (Shcheglovitov *et al.*, 2013; Nehme *et al.*, 2018). Representative traces are shown in **Fig. 2A–D.** I_Na_ density measurements are shown for −30 mV, the peak voltage for most neurons tested. P1 neurons had decreased transient I_Na_ density that approached significance (233 ± 39 pA/pF [n = 24] for controls vs. 140 ± 12 pA/pF [n = 24] for P1, p = .058, **Fig. 2I**). Persistent I_Na_ was measured at 50 ms after onset of a depolarizing step to −30 mV and normalized to peak I_Na_. Neurons from P1 and P2 had significantly higher percentages of persistent I_Na_ compared to controls (control: 3.6 ± 0.5 % [n = 24]; P1: 5.7 ± 0.6 % [n = 24], p = .012 ; P2: 6.81 ± 1.33 % [n = 10], p = .014, **Fig. 2J**). In contrast, persistent I_Na_ in P3 neurons were not significantly different from controls (P3: 3.61 ± 0.44 [n = 17], p = .985), suggesting an alternative mechanism.

**Figure 2.**
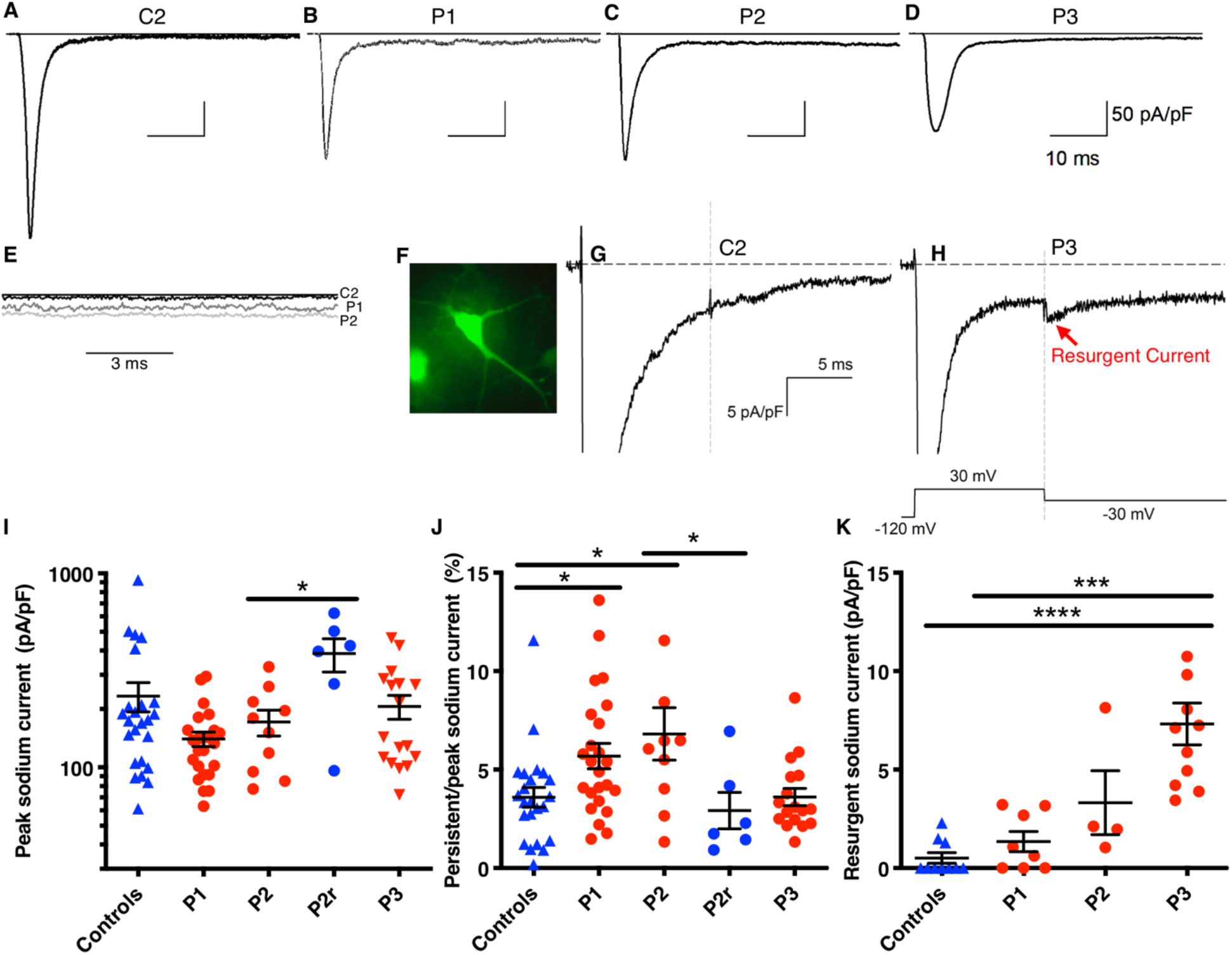
Increased persistent or resurgent INa in EIEE13 patient neurons. (**A-D**) Representative I_Na_ traces for control and patient neurons. (**E**) Persistent I_Na_ traces for C2, P1 and P2 are overlaid to demonstrate increased currents in patient neurons. (**F**) Representative epifluorescent image of a *Camk2a*-GFP labeled neuron from which whole-cell patch clamp recordings were made. (**G, H**) Resurgent I_Na_ recordings from C2 and P3, respectively. (**I, J**) Quantification of peak I_Na_ density at −30 mV membrane potential and persistent I_Na_ density 50 ms after depolarization divided by peak I_Na_, respectively. (**K**) Resurgent I_Na_ quantified after a 30 mV depolarization followed by −30 mV repolarization voltage step. Error bars are SEM. *<0.05, **<0.01,***<0.001, and **** <0.0001 by Kruskal-Wallis test with uncorrected Dunn’s post-test. For **I-J** controls N = 24 (C1 = 15, C2 = 9), P1 = 24 (Clone1 = 12, Clone2 = 12), P2 = 10 (Clone1 = 4, Clone2 = 6), P2r = 6, and P3 = 17 (Clone1 = 8, Clone2 = 5, Clone3 = 4). For *K* controls N = 10 (C1 = 6, C2 = 4), P1 = 8 (Clone1 = 3, Clone2 = 5), P2 = 4 (Clone1 = 1, Clone2 = 3), and P3 = 11 (Clone1 = 4, Clone2 = 4, Clone3 = 3).

Resurgent I_Na_ density was recorded using the voltage-step protocol shown in **Fig. 2H**. Representative traces are shown in **Fig. 2 G,H**. P3 neurons had significantly higher resurgent I_na_ density (7.3 pA/pF [n = 10]) compared to controls (0.5 pA/pF [n = 11], p<.0001) while P1 (1.3 pA/pF [n = 8], p = .375) and P2 neurons (3.3 pA/pF [n = 4], p = 0.0706) did not (**Fig. 2K**).

Voltage-dependent properties across the full I-V curve range for a subset of the recorded neurons were not significantly different between genotypes; however, a trend for decreased peak I_Na_ (G_max_) for P1 and P3 fits with the significant results found at −30 mV (**Supplemental Table 4**). Additionally, there are no differences in inactivation kinetics between controls and patient cells (**Supplemental Table 4**).

### Deletion of the P2 pathogenic allele rescues persistent I_Na_

An edited P2 iPSC line was generated using simultaneous CRISPR gene-editing and iPSC reprogramming. The “nickase” Cas9 variant, D10A, was used with two guide RNA sequences that flanked the patient variant. A silent mutation was engineered in the ssODN template to alter one of the PAM sequences to block further nickase breaks. One line with only the wild-type variant was identified; however, the silent mutation was not present (**Supplemental Fig. 1A**). Further long-range PCR amplified a ~500 bp deletion of the pathogenic allele (**Supplemental Fig. 3**).This P2 rescue line, which we call P2r, was differentiated using the same protocol as control and patient lines. Transient I_Na_ density at −30 mV was larger in the P2r line than in P2 neurons (P2r: 386 ± 75 pA/pF [n = 6]; P2: 171 ± 26 pA/pF [n = 10], p = .040, **Fig. 2I**), consistent with the observed trend for larger transient I_Na_ density values in neurons from controls compared to those from P1 and P2 (**Fig. 2I**). Because transient I_Na_ density in the CRISPR gene-edited line was similar to control levels, we concluded that hemizygous *SCN8A* expression was not deleterious. Importantly, the elevated percent persistent I_Na_ observed in P2 neurons was rescued in the gene edited line, P2r, with values similar to controls but significantly different from P2 (P2r: 2.9 ± 0.9 % [n = 6]; P2: 6.8 ± 1.3 % [n = 10], p = .015; controls: 3.6 ± 0.5 % [n = 24], p = .486, **Fig. 2J**).

### EIEE13 neurons have EADs and slowed AP repolarization

To investigate changes in neuronal excitability, we recorded spontaneous activity from *Camk2a*-GFP positive iPSC-derived neurons in current-clamp mode. We observed EADs in P3 neurons, similar to findings in the *Scn8a^N1768D/+^* EIEE13 mouse model (Lopez-Santiago *et al.*, 2017), but not in neurons from controls, P1 or P2 (**Fig. 3A, B** and data not shown). We evoked single APs with 1-ms current injections in P1 and P3 neurons. We focused primarily on these two patients because they had the most severe clinical phenotypes and represented both classes of I_Na_ abnormalities. Representative traces are depicted in **Fig. 3C.** P1 neurons showed prolonged repolarization and P3 neurons showed EADs (arrows). Many P1 and P3 neurons did not return to resting membrane potential levels within the duration of the AP recording. Because of this, standard ADP_50_ and ADP_80_ measurements could not be accurately determined. Instead, we used 3 time-points following the AP peak (5, 10, and 40 ms) to compare the 3 groups (**Fig. 3E–G**). The resulting peak amplitude values were similar between groups, demonstrating that these measures were appropriate (**Fig. 3D**). P1 neuronal APs were more depolarized than controls at 5 and 10 ms, respectively (Fig. 3E, F). APs in P3 neurons were more depolarized than controls at the 10 and 40 ms time points (Fig. 3F, G). Passive membrane properties, including resting membrane potential, input resistance, and capacitance, were similar between patients and controls (**Supplemental Table 3**).

**Figure 3.**
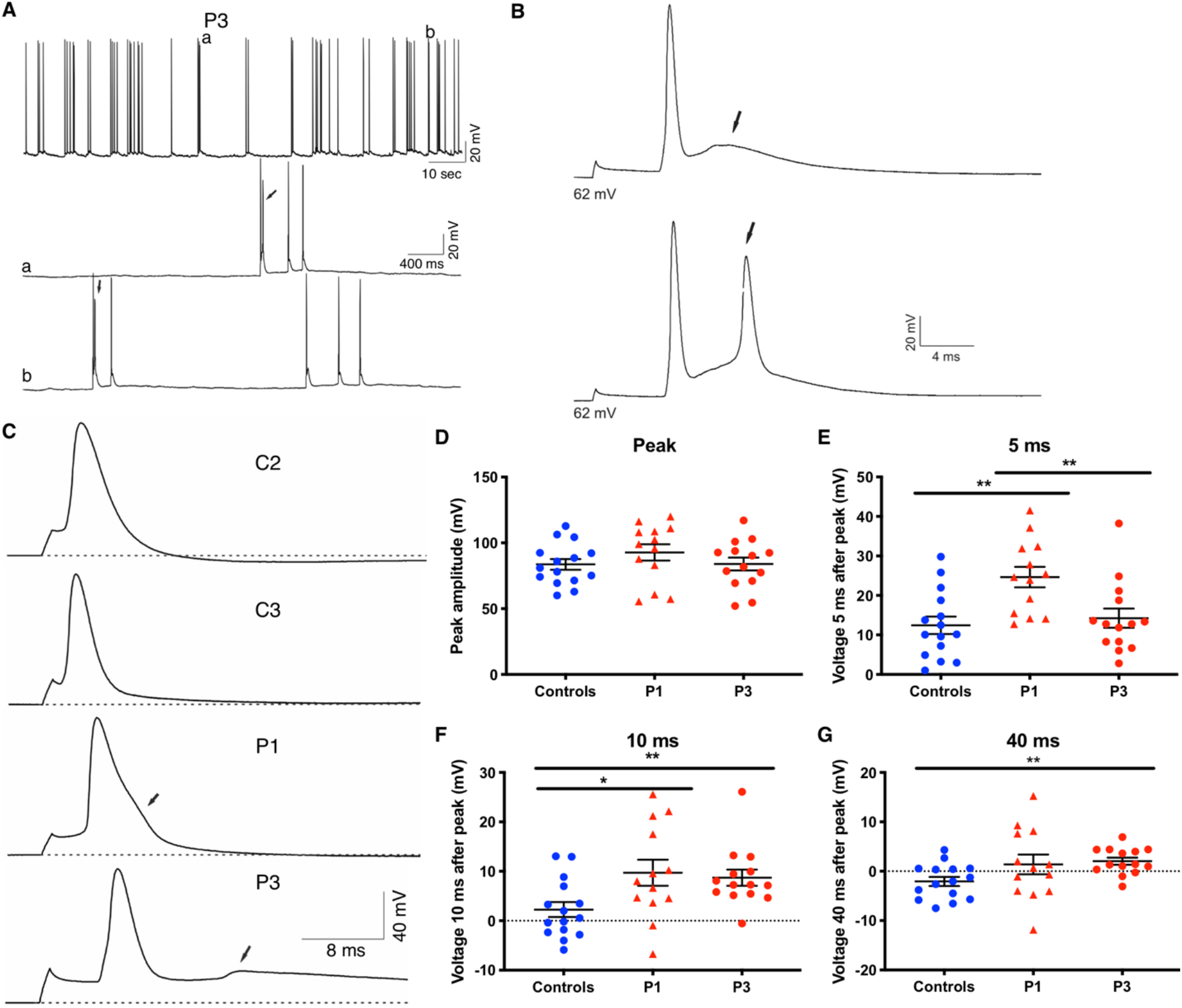
Impaired AP repolarization in EIEE13 patient iPSC-derived neurons. (**A**) Representative traces of spontaneous activity in P3 neurons. Arrows indicate instances of early afterdepolarization (EAD)-like waveforms. (**B**) Single spontaneous APs displayed with a higher resolution time scale show EAD-like waveforms in P3 neurons (arrows). (**C**) Examples of evoked AP firing for C2, C3, P1, and P3. Dotted lines denote the resting membrane potential (V_MR_). Arrows denote abnormalities in patient neuron AP repolarization. (**D**) The peak amplitude in mV as measured from the resting membrane potential to the peak potential. (**E-G**) Measurements of voltage difference from the resting membrane potential at 5, 10, and 40 ms after the initial evoked AP. Error bars represent standard error of the mean (SEM). * = p<0.05, ** = p<0.01 by Kruskal-Wallis test with uncorrected Dunn’s posttest.

### EIEE13 patient iPSC-derived neurons have reduced AIS length

Na_v_1.6, the protein encoded by *SCN8A*, primarily localizes to the neuronal AIS and nodes of Ranvier via interactions with its binding partner ankyrin-G (Gasser *et al.*, 2012). To determine whether neurons with EIEE13 variants display AIS alterations, we immunostained for ankyrin-G (Gutzmann *et al.*, 2014) as a proxy for the AIS in iPSC neurons. Nearly all neurons tested contained a single, well-defined AIS (**Fig. 4A–C**). We observed a significant decrease in AIS length in neurons derived from all three patient lines compared to controls (**Fig. 4D, E**). On average, the AIS length in patient neurons was decreased by 32% (38% for P1, 25% for P2, and 33% for P3) compared to the 3 controls.

**Figure 4.**
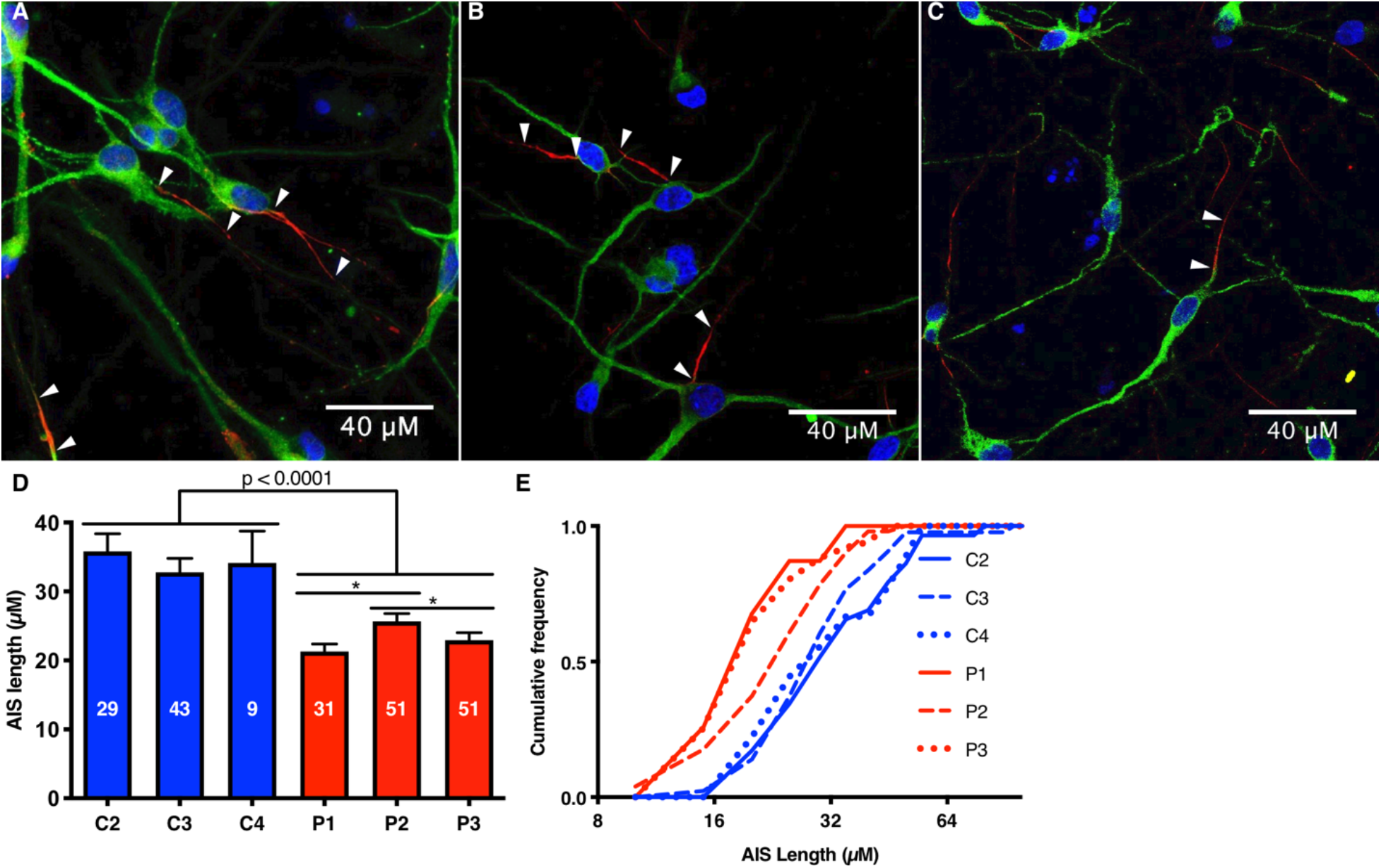
Decreased AIS length in EIEE13 patient-derived excitatory neurons. (**A-C**) Excitatory neurons from C1, P1, and P3, respectively, immunostained for ankyrin-G (red) and MAP2ab (green), with nuclear Hoechst labeling in blue. Arrowheads point to the start and end of each AIS from different neurons. (**D, E**) Quantification of AIS lengths for excitatory neurons from 3 controls and 3 patients plotted as bar graphs **D** with error bars as SEM, or as cumulative distributions **E** with 5 μM bin size. The N for each cell line is written in the bar graph in *D*. * denotes p<0.05 by Kruskal-Wallis test with uncorrected Dunn’s post-test.

### EIEE13 patient-derived iNeurons show differences in AP dynamics

We next used the iPSC neuron model for population analyses of neuronal activity and drug screening by MEA recordings. While the dual-SMAD differentiation technique paired with the *Camk2a* promoter-driven GFP reporter was useful for assessing individual neuron electrophysiology and AIS length, network activity assessed by MEA recordings was highly variable, possibly due to heterogeneous neuronal maturation and variability in the percentage of GABAergic neurons between cultures (data not shown). To reduce variability, we engineered iPSC lines to rapidly and homogeneously generate iNeurons by stably expressing dox-inducible *Ngn1* and *Ngn2*. As previously described, these cells quickly and efficiently differentiate into neurons after dox addition (Zhang *et al.*, 2013; Busskamp *et al.*, 2014; Lam *et al.*, 2017). *Ngn1/2* induction produces neurons expressing upper cortical layer markers, including *CUX1, CUX2,* and *BRN2*, but not deeper layer markers like *CTIP2* (Zhang *et al.*, 2013; Busskamp *et al.*, 2014; Nehme *et al.*, 2018). Stable insertion was successfully achieved for P1, P3, C2, and C3 lines using TALENs and homology arms targeting the *CLYBL* gene that is considered a “safe-harbor-like” locus in the human genome (Cerbini *et al.*, 2015). We first differentiated iNeurons (**Fig. 5B**) and measured evoked APs in single cells by patch clamp methods. APs from patient iNeurons showed reduced repolarization rates, similar to observations in the dual-SMAD neurons (controls: 17.1 ± 3.1 mV/ms [n = 37]; P1: 15.6 ± 3.8 mV/ms [n = 46], p = 0.028; P2: 14.6 ± 3.1 mV/ms [n = 29], p = 0.0038) (**Fig. 5A, F**). In addition, P1 and P3 cells showed increased AP amplitude half-widths compared to controls (controls: 3.08 ± 0.84 ms [n = 37]; P1: 3.56 ± 1.17 ms [n = 46], p = 0.027; P3: 3.48 ± 0.84 mV/ms [n = 29], p = 0.034) (**Fig. 5D**). Finally, P3 neurons exhibited reduced AP peak amplitude and rate of depolarization compared with controls (**Fig. 5C, E**).

**Figure 5.**
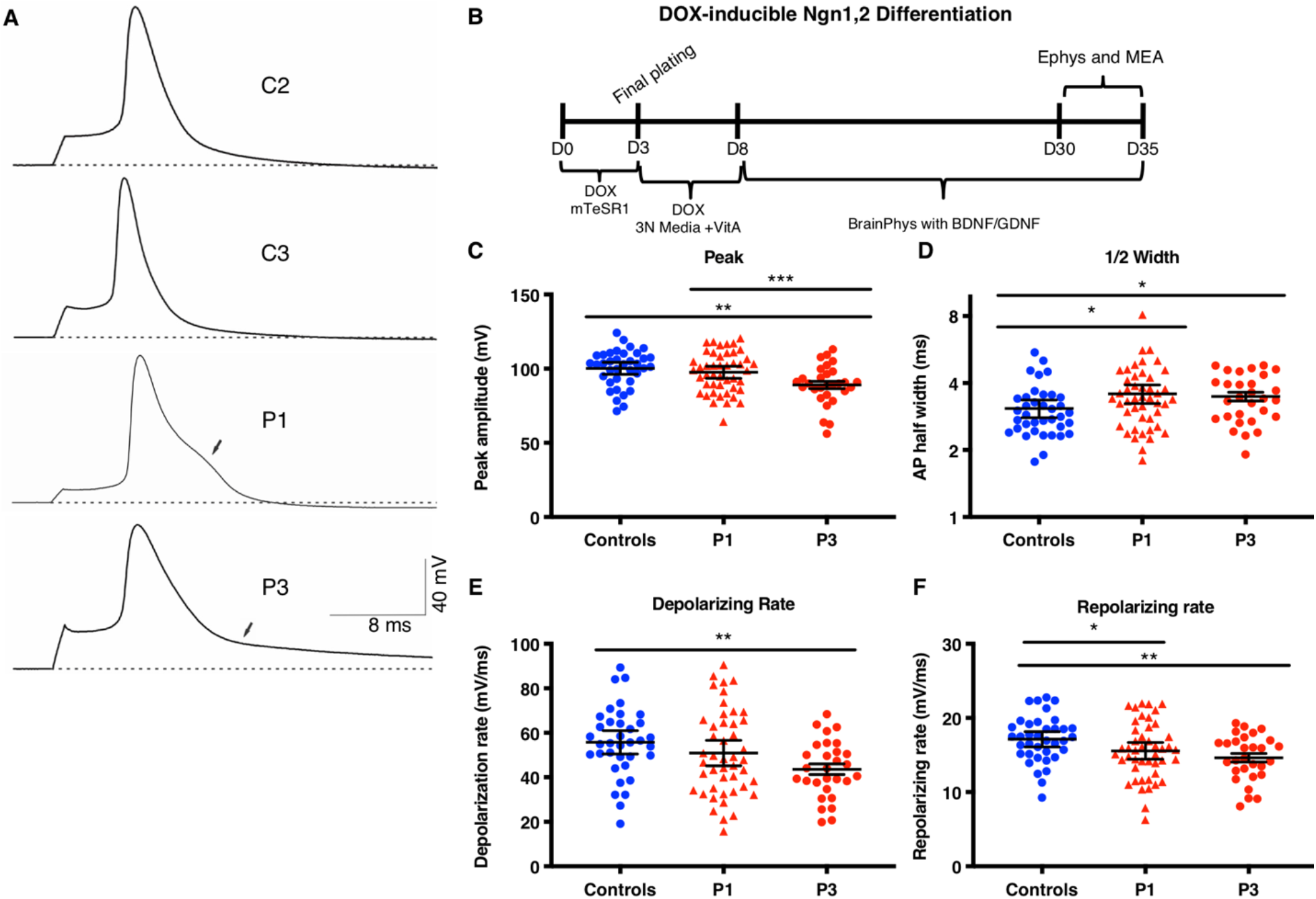
Altered AP repolarization in EIEE13 patient-derived iNeurons. (**A**) Representative traces of evoked AP firing for controls, P1, and P3. Dotted line denotes the resting membrane potential (V_MR_). Arrows denote abnormalities in patient neuron AP repolarization. (**B**) Schematic of iNeuron differentiation protocol from iPSC lines with stable cassette insertion in *CLYBL* safe-harbor-like locus. (**C-F**) Measurement of key AP characteristics including the peak amplitude in mV as measured from the resting membrane potential to the peak potential (**C**), AP width at half the peak height (**D**), and the rates of depolarization and repolarization (**E** and **F**, respectively). Error bars represent SEM. * = p<0.05, ** = p<0.01, *** = p<0.001 by Kruskal-Wallis test with uncorrected Dunn’s post-test.

### Increased burstiness of EIEE13 patient iNeurons

iNeurons were differentiated and plated onto 96-well MEA plates (detailed in Methods and **Fig. 5B**) and activity was measured for over a month. Individual raster plots and instantaneous firing frequency plots of neuronal activity on day 33 revealed that EIEE13 network activity, particularly that of P1 neurons, displayed periods of rapid network bursts that become shorter and more frequent with time, followed by a brief period of low activity (**Fig. 6A–C**). Both the increased burst activity and pattern of burst discharges in EIEE13 iNeurons are consistent with an epileptiform-like phenotype. After more than 4 weeks in culture (days 29-33), we found significant increases in measures of bursting activity in patient iNeurons compared to controls, as assessed by burst duration and the percentage of total spikes that were in network bursts for both P1 and P3 (**Fig. 6F, G, J, K**) (See **Methods and materials** for burst and network burst criteria). Another measure of burstiness, the coefficient of variation of interspike interval was elevated in P1 only (**Fig. 6E, I**) (Nawrot *et al.*, 2008). These measures of burstiness were not due to overall increases in firing, given that P1 tended to have lower overall activity compared with controls as measured by the weighted mean firing rate (MFR) and P3 was only slightly elevated (**Fig. 6D, H**).

**Figure 6.**
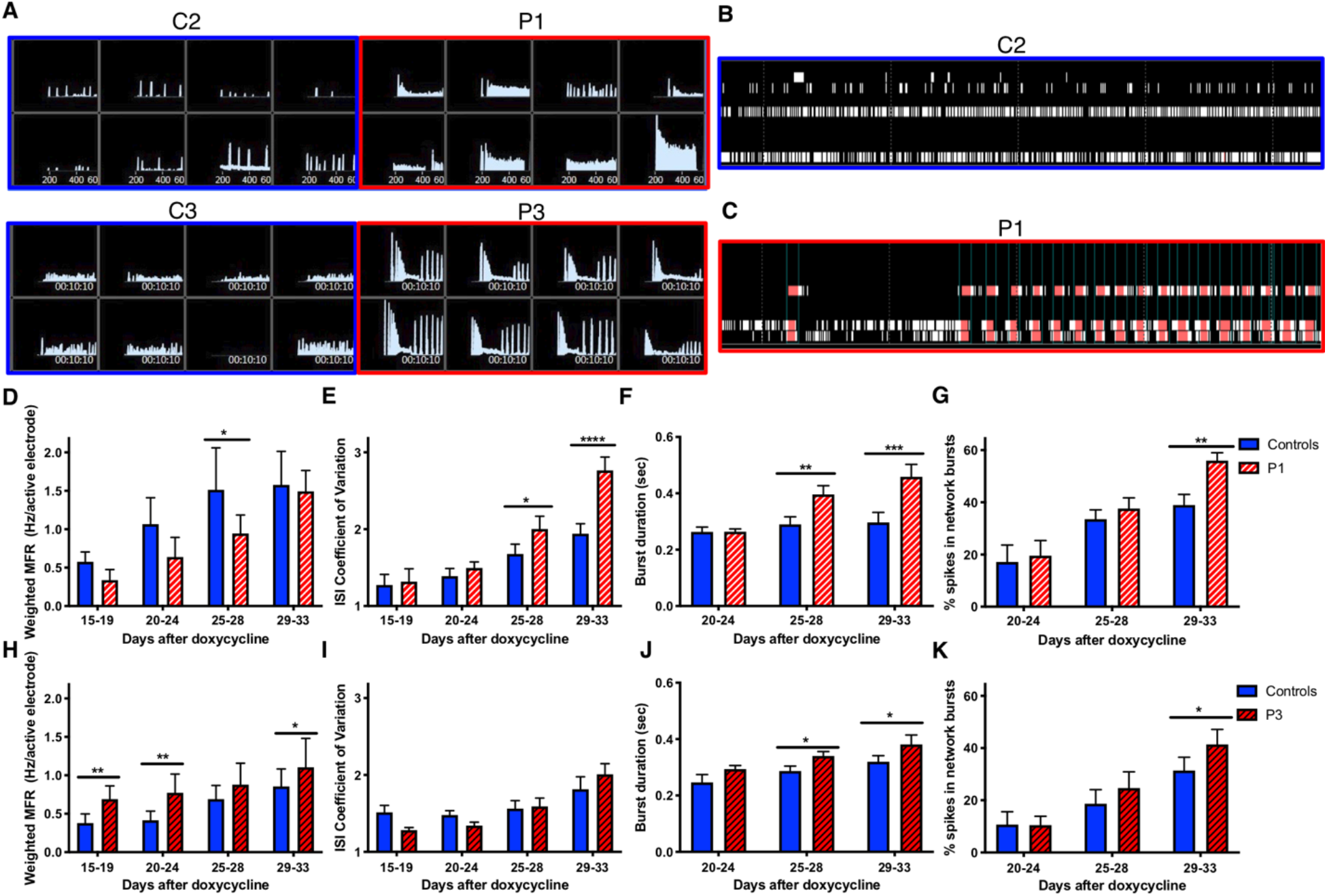
Increased burstiness in P1 and P3 iNeurons. (**A**) Control and patient iNeuron networks recorded by MEA 33 days after plating. Instantaneous firing frequencies over a 5-minute recording show a unique bursting patterns in patient lines compared to control. (**B, C**) Raster plots for the 8 electrodes in an individual well of a 96-well MEA plate for C2 and P1. (**D-K**) 8 paired MEA experiments were performed on 96-well MEA plates with 48 wells per line. Mean values were recorded for each line for each experiment. For some independent experiments, 2 identical plates were used. In this case the mean of the two plates was used for the individual data point. (**D,H**) The measurement of the mean firing frequency normalized to the number of active electrodes in a well. (**E, I**) The coefficient of variation for all interspike intervals (ISI). (**F, J**) Burst duration as defined by Axion software. (**G, K**) The % of total spikes in network bursts. Network bursts identification is defined in **Materials and methods**. (**D-G**) include both controls and P1: N = 8 controls (C2 = 6, C3 = 2), N = 8 P1. (**H-K**) include both controls and P3: N = 8 controls (C2 = 2, C3 = 6), N = 8 P3. * denotes p<.05, ** p<.01, *** p<.001, and **** p<.0001 by two-way ANOVA with two-stage linear step-up procedure post-test.

### Riluzole and phenytoin attenuate bursting phenotypes in EIEE13 patient iNeurons

We next tested the effects of the ASM phenytoin, which has been used with some success in EIEE13 patients to reduce seizures, as well as riluzole, a drug that has been shown to inhibit persistent and resurgent I_Na_ (Urbani and Belluzzi, 2000; Theile and Cummins, 2011). Riluzole also blocks EAD-like AP wave forms in the *Scn8a^N1768D/+^* EIEE13 mouse model without significant reduction of the rising phase of the AP (Lopez-Santiago *et al.*, 2017). Drug concentrations were chosen based on patient cerebrospinal fluid data from amyotrophic lateral sclerosis (ALS, riluzole) or epilepsy (phenytoin) patients (Groeneveld *et al.*, 2001; Rambeck *et al.*, 2006). Experiments included drug concentrations at a half-log above and a half-log below those values, respectively. In whole-cell patch clamp recordings, 3 μM riluzole completely and reversibly inhibited spontaneous AP firing of P3 neurons (**Fig. 7A**). In evoked firing experiments, riluzole did not inhibit the first AP but blocked all subsequent repetitive firing at nearly all current injections (data not shown). Riluzole also increased the voltage threshold for initiating a single AP (**Fig. 7B**) of P3 and control neurons.

**Figure 7.**
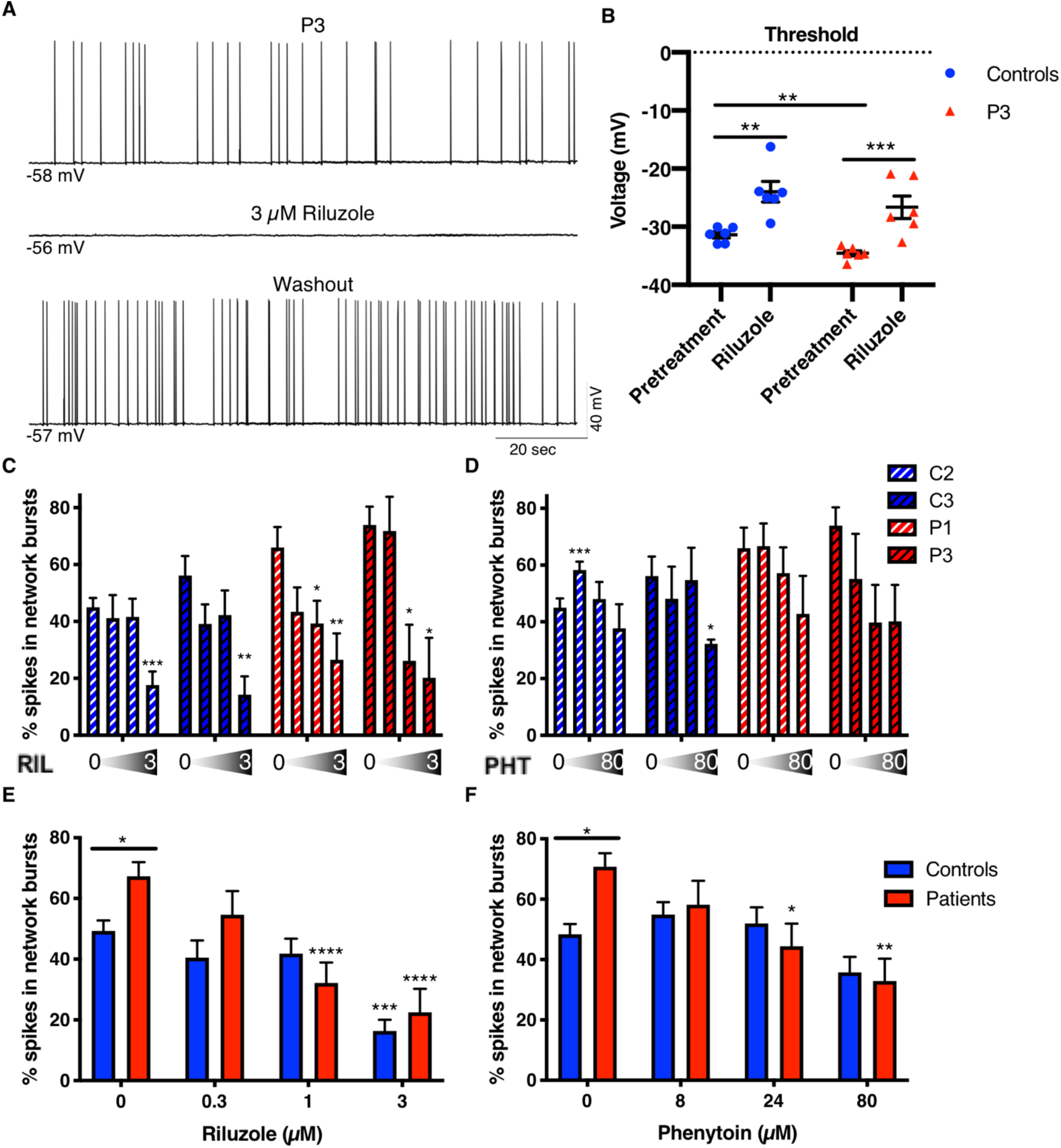
Phenytoin and riluzole suppress activity of EIEE13 patient iNeurons. (**A**) Patch-clamp recording of a spontaneously active P3 neuron before, during, and after washout of 3 μM riluzole. Riluzole inhibited all spontaneous activity but was completely reversible with washout. (**B**) The threshold voltage for single evoked APs was measured before and after riluzole exposure. For both controls and P3, riluzole raised the AP threshold to more depolarized potentials. (**C, D**) Changes in % spikes in network bursts in patient and control iNeurons in response to increasing concentrations of riluzole (**C**; 0, 0.3, 1, and 3 μM) and phenytoin (**D**; 0, 8, 24, and 80 μM). N = 8 independent experiments for C2 and P1, respectively. N = 5 independent experiments for C3 and P3, respectively. Within group comparisons for drug dosage effect compared to vehicle (DMSO) were performed using a one-way ANOVA. (**E, F**) Grouped control and patient data from **C** and **D**, respectively. N = 13 for each group. Two-way unpaired ANOVA (due to missing values) was performed. All post-hoc analyses used the two-stage linear step-up procedure post-test. *<0.05, **<0.01, ***<0.001, and ****<0.0001.

Acute effects of the drugs were determined by MEA recordings following a 5-minute equilibration time of the drugs in the culture wells. Since the percentage of spikes in network bursts was the most reliable excitability difference for the two patient lines compared to controls (**Fig. 6G, K**), we compared the effects of riluzole and phenytoin on this parameter as well as on the MFR. We found that 1 μM riluzole significantly decreased the percentage of spikes in network bursts in patient, but not control, iNeuron cultures (**Fig. 7C, E**). A similar, patient-specific effect was seen with 24 μM phenytoin (**Fig. 7D, F**). Two-way ANOVA analysis from vehicle to therapeutic concentration (1 μM for riluzole and 24 μM for phenytoin) had a significant genotype by treatment interaction (riluzole: p = 0.036, phenytoin: p = 0.035) meaning that the cells responded to both drugs in a genotype-dependent manner. When we plotted the MFR normalized to the pretreatment MFR, we observed a steady decrease in activity with increasing dosage but no differences in the effect of either drug between patient and control (**Supplemental Fig. 4**). These results indicate an increased sensitivity of bursting activity in patient iNeurons for both drugs compared to controls, with no genotype-related differential drug effects on overall activity. Notably, over the concentration range tested, riluzole had a greater inhibitory effect on both percentage of spikes in network bursts (**Fig. 7**) and overall MFR (**Supplemental Fig. 4**) than phenytoin.

## Discussion

Our results demonstrate the utility of iPSC-derived neurons to identify variant-specific I_Na_ abnormalities in EIEE13 patients. Of three patients with distinct *SCN8A* variants studied, cortical-like neurons from two showed increased proportions of persistent I_Na_ and one showed increased resurgent I_Na_. All EIEE13 patient neurons had prolonged AP repolarization and decreased AIS lengths. iNeurons generated with the dox-inducible *Ngn1,2* system recapitulated the prolonged AP repolarization observed in patient neurons and demonstrated increased network bursting measured by MEA recordings. These recordings provided a platform for testing the VGSC-selective drugs, phenytoin and riluzole. Riluzole administration produced similar effects as phenytoin, one of the most widely used therapies for EIEE13 (Boerma *et al.*, 2016).Both drugs reduced network bursting activity at pharmacologically relevant concentrations with some specificity for the disease variant neuronal networks, suggesting that further study of riluzole or riluzole derivatives as an EIEE13 therapy is warranted.

The increase in persistent I_Na_ that we identified in human iPSC-derived neurons is consistent with previous studies in other model systems. Elevated persistent I_Na_ was reported in the first identified EIEE13 disease variant (N1768>D) modeled in transduced hippocampal neurons (Veeramah *et al.*, 2012). A mouse model of the N1768>D variant displayed increased persistent I_Na_, slowed repolarization, and EAD-like AP waveforms in CA1 hippocampal neurons (Lopez-Santiago *et al.*, 2017). Immortalized cells transiently expressing *SCN8A* cDNA with the N1768>D variant exhibited both increased persistent *and* resurgent I_Na_ (Patel *et al.*, 2016), suggesting that the effect of this mutation on I_Na_ may be cell type dependent. Here, we found increased persistent I_Na_ in patient-derived neuronal cultures expressing two different *SCN8A* variants. However, unlike P1 (R1872>L) and P2 (V1592>L) neurons, P3 neurons (N1759>S) showed unaltered levels of persistent I_Na_ but, instead, showed elevated resurgent I_Na_ density. Neurons from P3, but not from P1 or P2, also exhibited EAD-like waveforms during spontaneous firing. A recent study of variants in *SCN9A* linked to small fiber neuropathy described increased resurgent I_Na_, broader APs, and EADs (Xiao *et al.*, 2019), similar changes as described here for P3 neurons. These data suggest that elevated resurgent I_Na_ may be an underlying mechanism for the development of EAD-like waveforms in some types of mutant neurons.

In addition to the clear differences in I_Na_ recorded between patient and control neurons, our hemizygous P2r rescue experiment provides an isogenic control in the absence of the disease variant to demonstrate that the observed increase in persistent I_Na_ in P2 neurons is due to the *SCN8A de novo* variant, rather than to other genetic differences. While haploinsufficiency of some VGSC genes can lead to epilepsy (e.g. *SCN1A* in Dravet syndrome), heterozygous *SCN8A* loss-of-function in mice results in no overt phenotype, except subtle absence seizures (Papale *et al.*, 2009). In fact, the *Scn8a^med-jo/+^* mouse crossed with a Dravet model (*Scn1a^+/-^)* resulted in rescue of induced seizure threshold to wild-type levels (Martin *et al.*, 2007). Taken together, this information and the observation that transient I_Na_ was not reduced in P2r neurons suggest that this line is an appropriate rescue for P2 neurons.

The observed reduction in AIS length in EIEE13 patient neurons was unexpected. Others have proposed that AIS shortening is a compensatory mechanism to counter hyperexcitability, and animal models have shown AIS shortening in cortical neurons after TBI or stroke (Baalman *et al.*, 2013; Hinman *et al.*, 2013; Evans *et al.*, 2015). AIS length is also known to vary with developmental timing and input in the visual cortex (Gutzmann *et al.*, 2014). More work is necessary to determine whether AIS shortening in EIEE13 neurons reflects a compensatory attempt to decrease hyperexcitability or instead relates to altered development. Interestingly, in terms of the latter possibility, we did not observe AIS shortening in iNeurons (data not shown), which may lose developmental phenotypes observed with small molecule differentiation protocols that more faithfully recapitulate developmental events (Schafer *et al.*, 2019).

MEA recordings demonstrated increased burstiness in P1 and P3 iNeuron cultures. We used iNeurons to decrease variability by forcing synchronized differentiation into pure excitatory cortical-like neurons. While this approach decreased the variability in our experiments considerably, purely excitatory cultures exhibit spontaneous network burst firing, potentially an epileptiform-like event, under basal conditions (Matsuda *et al.*, 2018). While both control and patient iNeuron networks exhibited these events, the EIEE13 patient iNeuron networks had greater “burstiness” than controls as measured by longer burst durations and greater percentage of spikes in network bursts. P1 neurons also displayed a greater interspike interval coefficient of variation than controls (**Fig. 6F**). These measures suggest increased activity within network bursts, likely due to the cell-intrinsic *SCN8A* gain-of-function mechanisms we identified using single cell electrophysiological analyses. These differences are similar to changes in network bursts observed following the administration of chemoconvulsants such as bicuculline, pentylentetrazole, and 4-aminopyridine (Odawara *et al.*, 2016). A recent publication investigating Kleefstra syndrome using iNeurons on MEAs identified disease specific alterations using similar outcomes measures, including burst duration, % of spikes out of bursts, and inter-burst-interval coefficient of variation (Frega *et al.*, 2019). Importantly, EIEE13 patient cultures demonstrated “burst of bursts” events with high-frequency network bursting that did not occur in controls. These events were often followed by reduced activity, resulting in no significant differences in burst frequency calculated over the 5-minute recording period. The reproducible differences in bursting parameters we observed between EIEE13 patient and control neuronal networks allowed us to begin assessing the effects of anti-epileptic drugs.

Currently, the VGSC blocking ASMs, including high-dose phenytoin, are the most effective and commonly used therapies for patients with *SCN8A* EIEE13 variants (Boerma *et al.*, 2016; Gardella *et al.*, 2018). However, these drugs frequently cause adverse effects and most EIEE13 patients continue to have frequent seizures despite the use of multiple ASMs (Braakman *et al.*, 2017). We chose to examine a VGSC-selective inhibitor not currently used for epilepsy, riluzole, for its potential to more specifically attenuate persistent and resurgent I_Na_ than phenytoin. Riluzole is used for the treatment of ALS and is known to inhibit I_Na_ by binding to and stabilizing the inactive state of VGSC α subunits (Song *et al.*, 1997). Riluzole has also been shown to preferentially inhibit both persistent and resurgent I_Na_ (Urbani and Belluzzi, 2000; Spadoni *et al.*, 2002; Theile and Cummins, 2011; Xie *et al.*, 2011). Even though riluzole has been used to effectively inhibit seizures in mice, rats, and baboons, it has not been used to treat epilepsy patients (Mizoule *et al.*, 1985; Romettino *et al.*, 1991; Doble, 1996; De Sarro *et al.*, 2000; Yoshida *et al.*, 2001). Although riluzole affects a wide array of channels and receptors, at the concentrations found in patients and tested in our MEA recordings (<10 μM), riluzole is thought to selectively inhibit persistent I_Na_ and potentiate calcium-dependent potassium currents (Bellingham, 2011). Application of either phenytoin or riluzole in concentrations reflecting patient CSF concentrations identified in the literature resulted in a variant-specific reduction in burstiness parameters to control levels (Groeneveld *et al.*, 2001; Rambeck *et al.*, 2006). Thus, our drug-testing platform is informative at treatment-relevant concentrations.

Patients 1, 3, and 4 experienced medically refractory seizures despite using standard VGSC blocking (including phenytoin, carbamazepine, oxcarbazepine, lamotrigine, etc) and other ASMs. The mutation position in the Na_V_1.6 channel may be, in part, why VGSC-blocking drugs were ineffective in patients 3 and 4 (**Table 1).** Previous work showed that the *SCN2A/*Nav1.2 variants N1769>A and V1767>A (among others on segment IVS6) altered the effectiveness of etidocaine on I_Na_ (Ragsdale *et al.*, 1994). The equivalent residues in Nav1.6 are N1759 (mutated in P3) and V1757 (mutated in P4). Neither patient had adequate seizure relief with typical ASMs – phenytoin, lamotrigine, and carbamazepine – that are thought to interact at this same site (Lipkind and Fozzard, 2010). Riluzole, on the other hand, has a theoretical binding pocket interacting with residues 1787, 1801, and 1845 of Na_V_1.6 (Bello *et al.*, 2012). These observations demonstrate that further studies are needed to assess the efficacy and safety of riluzole treatment in EIEE13. Although it should be noted that riluzole has been well tolerated by children with spinal muscular atrophy and obsessive-compulsive disorder (Russman *et al.*, 2003; Grant *et al.*, 2014). Importantly, the findings by our group in patient-derived neurons and others in heterologous systems/cultured neurons (Barker *et al.*, 2016; Wagnon *et al.*, 2016) of mutation-specific gain-of-function effects in EIEE13 suggests that the ASM choice may be variant-specific. The patient-derived neuronal models that we describe offer a useful approach for this type of precision epilepsy therapy.

## Acknowledgements

We thank the clinicians, parents, and patients who provided the skin biopsies and clinical information for this study. We also thank Dr. Michael Ward and the Inherited Neurodegenerative Diseases Unit at NINDS for the gift of the pUCM-CLYBL-Ngn1&2 plasmid.

## References

Baalman KL, Cotton RJ, Rasband SN, Rasband MN. Blast wave exposure impairs memory and decreases axon initial segment length. Journal of neurotrauma 2013; 30(9): 741–51.

Barker BS, Ottolini M, Wagnon JL, Hollander RM, Meisler MH, Patel MK. The SCN8A encephalopathy mutation p. Ile1327Val displays elevated sensitivity to the anticonvulsant phenytoin. Epilepsia 2016; 57(9): 1458–66.

Bellingham MC. A review of the neural mechanisms of action and clinical efficiency of riluzole in treating amyotrophic lateral sclerosis: what have we learned in the last decade? CNS neuroscience & therapeutics 2011; 17(1): 4–31.

Benjamini Y, Krieger AM, Yekutieli D. Adaptive linear step-up procedures that control the false discovery rate. Biometrika 2006; 93(3): 491–507.

Boerma RS, Braun KP, van de Broek MP, van Berkestijn FM, Swinkels ME, Hagebeuk EO, et al. Remarkable phenytoin sensitivity in 4 children with SCN8A-related epilepsy: a molecular neuropharmacological approach. Neurotherapeutics 2016; 13(1): 192–7.

Braakman HM, Verhoeven JS, Erasmus CE, Haaxma CA, Willemsen MH, Schelhaas HJ. Phenytoin as a last-resort treatment in SCN 8A encephalopathy. Epilepsia open 2017; 2(3): 343–4.

Bunton-Stasyshyn RKA, Wagnon JL, Wengert ER, Barker BS, Faulkner A, Wagley PK, et al. Prominent role of forebrain excitatory neurons in SCN8A encephalopathy. Brain 2019; 142(2): 362–75.

Busskamp V, Lewis NE, Guye P, Ng AH, Shipman SL, Byrne SM, et al. Rapid neurogenesis through transcriptional activation in human stem cells. Molecular systems biology 2014; 10(11): 760.

Cerbini T, Funahashi R, Luo Y, Liu C, Park K, Rao M, et al. Transcription activator-like effector nuclease (TALEN)-mediated CLYBL targeting enables enhanced transgene expression and one-step generation of dual reporter human induced pluripotent stem cell (iPSC) and neural stem cell (NSC) lines. PLoS One 2015; 10(1): e0116032.

De Sarro G, Siniscalchi A, Ferreri G, Gallelli L, De Sarro A. NMDA and AMPA/kainate receptors are involved in the anticonvulsant activity of riluzole in DBA/2 mice. European journal of pharmacology 2000; 408(1): 25–34.

Doble A. The pharmacology and mechanism of action of riluzole. Neurology 1996; 47(6 Suppl 4): 233S–41S.

Epi4K, Investigators E. De novo mutations in the classic epileptic encephalopathies. Nature 2013; 501(7466): 217.

Evans MD, Dumitrescu AS, Kruijssen DL, Taylor SE, Grubb MS. Rapid modulation of axon initial segment length influences repetitive spike firing. Cell reports 2015; 13(6): 1233–45.

Frega M, Linda K, Keller JM, Gümüş-Akay G, Mossink B, van Rhijn J-R, et al. Neuronal network dysfunction in a model for Kleefstra syndrome mediated by enhanced NMDAR signaling. Nature communications 2019; 10(1): 1–15.

Gardella E, Marini C, Trivisano M, Fitzgerald MP, Alber M, Howell KB, et al. The phenotype of SCN8A developmental and epileptic encephalopathy. Neurology 2018; 91(12): e1112–e24.

Gasser A, Ho TS-Y, Cheng X, Chang K-J, Waxman SG, Rasband MN, et al. An ankyrinG-binding motif is necessary and sufficient for targeting Nav1. 6 sodium channels to axon initial segments and nodes of Ranvier. Journal of Neuroscience 2012; 32(21): 7232–43.

Grant PJ, Joseph LA, Farmer CA, Luckenbaugh DA, Lougee LC, Zarate Jr CA, et al. 12-week, placebo-controlled trial of add-on riluzole in the treatment of childhood-onset obsessive– compulsive disorder. Neuropsychopharmacology 2014; 39(6): 1453.

Groeneveld G, Van Kan H, Toraño JS, Veldink J, Guchelaar H-J, Wokke J, et al. Inter-and intraindividual variability of riluzole serum concentrations in patients with ALS. Journal of the neurological sciences 2001; 191(1): 121–5.

Gutzmann A, Ergül N, Grossmann R, Schultz C, Wahle P, Engelhardt M. A period of structural plasticity at the axon initial segment in developing visual cortex. Frontiers in neuroanatomy 2014; 8.

Hinman JD, Rasband MN, Carmichael ST. Remodeling of the axon initial segment after focal cortical and white matter stroke. Stroke 2013; 44(1): 182–9.

Kong W, Zhang Y, Gao Y, Liu X, Gao K, Xie H, et al. SCN8A mutations in Chinese children with early onset epilepsy and intellectual disability. Epilepsia 2015; 56(3): 431–8.

Lam RS, Töpfer FM, Wood PG, Busskamp V, Bamberg E. Functional maturation of human stem cell-derived neurons in long-term cultures. PloS one 2017; 12(1): e0169506.

Larsen J, Carvill GL, Gardella E, Kluger G, Schmiedel G, Barisic N, et al. The phenotypic spectrum of SCN8A encephalopathy. Neurology 2015; 84(5): 480–9.

Lipkind GM, Fozzard HA. Molecular model of anticonvulsant drug binding to the voltage-gated sodium channel inner pore. Molecular pharmacology 2010: mol. 110.064683.

Liu Y, Lopez-Santiago LF, Yuan Y, Jones JM, Zhang H, O'malley HA, et al. Dravet syndrome patient-derived neurons suggest a novel epilepsy mechanism. Annals of neurology 2013; 74(1): 128–39.

Lopez-Santiago LF, Yuan Y, Wagnon JL, Hull JM, Frasier CR, O’Malley HA, et al. Neuronal hyperexcitability in a mouse model of SCN8A epileptic encephalopathy. Proceedings of the National Academy of Sciences 2017; 114(9): 2383–8.

Maroof AM, Keros S, Tyson JA, Ying S-W, Ganat YM, Merkle FT, et al. Directed differentiation and functional maturation of cortical interneurons from human embryonic stem cells. Cell stem cell 2013; 12(5): 559–72.

Martin MS, Tang B, Papale LA, Yu FH, Catterall WA, Escayg A. The voltage-gated sodium channel Scn8a is a genetic modifier of severe myoclonic epilepsy of infancy. Human molecular genetics 2007; 16(23): 2892–9.

Matsuda N, Odawara A, Katoh H, Okuyama N, Yokoi R, Suzuki I. Detection of synchronized burst firing in cultured human induced pluripotent stem cell-derived neurons using a 4-step method. Biochemical and biophysical research communications 2018; 497(2): 612–8.

Mizoule J, Meldrum B, Mazadier M, Croucher M, Ollat C, Uzan A, et al. 2-Amino-6-trifluoromethoxy benzothiazole, a possible antagonist of excitatory amino acid neurotransmission—I: anticonvulsant properties. Neuropharmacology 1985; 24(8): 767–73.

Nawrot MP, Boucsein C, Molina VR, Riehle A, Aertsen A, Rotter S. Measurement of variability dynamics in cortical spike trains. Journal of neuroscience methods 2008; 169(2): 374–90.

Neely MD, Litt MJ, Tidball AM, Li GG, Aboud AA, Hopkins CR, et al. DMH1, a highly selective small molecule BMP inhibitor promotes neurogenesis of hiPSCs: comparison of PAX6 and SOX1 expression during neural induction. ACS chemical neuroscience 2012; 3(6): 482–91.

Nehme R, Zuccaro E, Ghosh SD, Li C, Sherwood JL, Pietilainen O, et al. Combining NGN2 Programming with Developmental Patterning Generates Human Excitatory Neurons with NMDAR-Mediated Synaptic Transmission. Cell reports 2018; 23(8): 2509–23.

Odawara A, Katoh H, Matsuda N, Suzuki I. Physiological maturation and drug responses of human induced pluripotent stem cell-derived cortical neuronal networks in long-term culture. Scientific reports 2016; 6: 26181.

Ohba C, Kato M, Takahashi S, Lerman-Sagie T, Lev D, Terashima H, et al. Early onset epileptic encephalopathy caused by de novo SCN8A mutations. Epilepsia 2014; 55(7): 994–1000.

Ottolini M, Barker BS, Gaykema RP, Meisler MH, Patel MK. Aberrant sodium channel currents and hyperexcitability of medial entorhinal cortex neurons in a mouse model of SCN8A encephalopathy. Journal of Neuroscience 2017: 2709–16.

Papale LA, Beyer B, Jones JM, Sharkey LM, Tufik S, Epstein M, et al. Heterozygous mutations of the voltage-gated sodium channel SCN8A are associated with spike-wave discharges and absence epilepsy in mice. Human molecular genetics 2009; 18(9): 1633–41.

Patel RR, Barbosa C, Brustovetsky T, Brustovetsky N, Cummins TR. Aberrant epilepsy-associated mutant Nav1. 6 sodium channel activity can be targeted with cannabidiol. Brain 2016; 139(8): 2164–81.

Perreault ML, Fan T, Alijaniaram M, O'Dowd BF, George SR. Dopamine D1–D2 receptor heteromer in dual phenotype GABA/glutamate-coexpressing striatal medium spiny neurons: regulation of BDNF, GAD67 and VGLUT1/2. PloS one 2012; 7(3): e33348.

Ragsdale DS, McPhee JC, Scheuer T, Catterall WA. Molecular determinants of state-dependent block of Na+ channels by local anesthetics. Science 1994; 265(5179): 1724–8.

Rambeck B, Jürgens UH, May TW, Pannek HW, Behne F, Ebner A, et al. Comparison of brain extracellular fluid, brain tissue, cerebrospinal fluid, and serum concentrations of antiepileptic drugs measured intraoperatively in patients with intractable epilepsy. Epilepsia 2006; 47(4): 681–94.

Romettino S, Lazdunski M, Gottesmann C. Anticonvulsant and sleep-waking influences of riluzole in a rat model of absence epilepsy. European journal of pharmacology 1991; 199(3): 371–3.

Root DH, Zhang S, Barker DJ, Miranda-Barrientos J, Liu B, Wang H-L, et al. Selective Brain Distribution and Distinctive Synaptic Architecture of Dual Glutamatergic-GABAergic Neurons. Cell Reports 2018; 23(12): 3465–79.

Russman BS, Iannaccone ST, Samaha FJ. A phase 1 trial of riluzole in spinal muscular atrophy. Archives of neurology 2003; 60(11): 1601–3.

Schafer ST, Paquola AC, Stern S, Gosselin D, Ku M, Pena M, et al. Pathological priming causes developmental gene network heterochronicity in autistic subject-derived neurons. Nature neuroscience 2019; 22(2): 243.

Shcheglovitov A, Shcheglovitova O, Yazawa M, Portmann T, Shu R, Sebastiano V, et al. SHANK3 and IGF1 restore synaptic deficits in neurons from 22q13 deletion syndrome patients. Nature 2013; 503(7475): 267.

Shi Y, Kirwan P, Smith J, Robinson HP, Livesey FJ. Human cerebral cortex development from pluripotent stem cells to functional excitatory synapses. Nature Neuroscience 2012; 15(3): 477–86.

Song J-H, Huang C-S, Nagata K, Yeh JZ, Narahashi T. Differential action of riluzole on tetrodotoxin-sensitive and tetrodotoxin-resistant sodium channels. Journal of Pharmacology and Experimental Therapeutics 1997; 282(2): 707–14.

Spadoni F, Hainsworth AH, Mercuri NB, Caputi L, Martella G, Lavaroni F, et al. Lamotrigine derivatives and riluzole inhibit INa, P in cortical neurons. Neuroreport 2002; 13(9): 1167–70.

Theile JW, Cummins TR. Inhibition of Navβ4 peptide-mediated resurgent sodium currents in Nav1. 7 channels by carbamazepine, riluzole, and anandamide. Molecular pharmacology 2011; 80(4): 724–34.

Tidball AM, Dang LT, Glenn TW, Kilbane EG, Klarr DJ, Margolis JL, et al. Rapid generation of human genetic loss-of-function iPSC lines by simultaneous reprogramming and gene editing. Stem cell reports 2017; 9(3): 725–31.

Tidball AM, Neely MD, Chamberlin R, Aboud AA, Kumar KK, Han B, et al. Genomic instability associated with p53 knockdown in the generation of huntington’s disease human induced pluripotent stem cells. PloS one 2016; 11(3): e0150372.

Tidball AM, Swaminathan P, Dang LT, Parent J. Generating loss-of-function iPSC lines with combined CRISPR indel formation and reprogramming from human fibroblasts. Bio Protoc 2018; 8.

Urbani A, Belluzzi O. Riluzole inhibits the persistent sodium current in mammalian CNS neurons. European Journal of Neuroscience 2000; 12(10): 3567–74.

Veeramah KR, O'Brien JE, Meisler MH, Cheng X, Dib-Hajj SD, Waxman SG, et al. De novo pathogenic SCN8A mutation identified by whole-genome sequencing of a family quartet affected by infantile epileptic encephalopathy and SUDEP. The American Journal of Human Genetics 2012; 90(3): 502–10.

Wagnon JL, Barker BS, Hounshell JA, Haaxma CA, Shealy A, Moss T, et al. Pathogenic mechanism of recurrent mutations of SCN8A in epileptic encephalopathy. Annals of clinical and translational neurology 2016; 3(2): 114–23.

Wagnon JL, Korn MJ, Parent R, Tarpey TA, Jones JM, Hammer MF, et al. Convulsive seizures and SUDEP in a mouse model of SCN8A epileptic encephalopathy. Human molecular genetics 2014; 24(2): 506–15.

Wagnon JL, Meisler MH. Recurrent and non-recurrent mutations of SCN8A in epileptic encephalopathy. Frontiers in neurology 2015; 6.

Xiao Y, Barbosa C, Pei Z, Xie W, Strong JA, Zhang J-M, et al. Increased resurgent sodium currents in Nav1. 8 contribute to nociceptive sensory neuron hyperexcitability associated with peripheral neuropathies. Journal of Neuroscience 2019; 39(8): 1539–50.

Xie R-G, Zheng D-W, Xing J-L, Zhang X-J, Song Y, Xie Y-B, et al. Blockade of persistent sodium currents contributes to the riluzole-induced inhibition of spontaneous activity and oscillations in injured DRG neurons. PLoS One 2011; 6(4): e18681.

Yau KW, van Beuningen SF, Cunha-Ferreira I, Cloin BM, van Battum EY, Will L, et al. Microtubule minus-end binding protein CAMSAP2 controls axon specification and dendrite development. Neuron 2014; 82(5): 1058–73.

Yoshida M, Noguchi E, Tsuru N, Ohkoshi N. Effect of riluzole on the acquisition and expression of amygdala kindling. Epilepsy research 2001; 46(2): 101–9.

Zhang Y, Pak C, Han Y, Ahlenius H, Zhang Z, Chanda S, et al. Rapid single-step induction of functional neurons from human pluripotent stem cells. Neuron 2013; 78(5): 785–98.

