## Supplemental material for "Variant-specific changes in persistent or resurgent Na^+^ current in *SCN8A*-EIEE13 iPSC-derived neurons"

#### Extended methods and materials

##### Induced pluripotent stem cell reprogramming

Skin punch biopsies were obtained from 3 EIEE13 patients (P1-3) and two healthy controls (C1 and C4) without known genetic disorders with consent under a protocol approved by the Institutional Review Board of Michigan Medicine. The biopsies were cut into several pieces and allowed to attach to several wells of a 6-well cell culture plate in Dulbecco's Modified Eagle Media (DMEM) with 10% fetal bovine serum (FBS), non-essential amino acids, and penicillin/streptomycin. After ~2 weeks, skin punches were removed and the attached dividing fibroblasts on the plate were passaged with trypsin (0.25%, Gibco). Media was exchanged every 2-3 days. The cells were passaged whenever they neared confluency. Fibroblasts ($1x10^5$) between passages 5-10 were electroporated in Neon electroporation kit reagent R premixed with reprogramming plasmids (1 µg each of pCXLE-hOCT3/4-p53shRNA, pCXLE-hUL, and pCXLE-hSK ) using the Neon Transfection system (ThermoFisher) with 3 pulses of 1650 V for 10 ms each as previously described (Okita *et al.*, 2011). Fibroblasts ($4-20 \times 10^3$) were then plated into each well of a Matrigel-coated 6-well plate in fibroblast growth medium. The fibroblast growth medium was changed the following day. Three days after electroporation, medium was changed to TeSR-E7 (Stem Cell Technologies) and replaced daily. iPSC clones were picked between days 17-21 onto Matrigel-coated dishes in mTeSR1 medium, which was exchanged daily. When cultures reached ~40% confluency, they were passaged using dispase and replated at a 1:4-1:8 dilution as small clumps (Thermo Fisher). Two additional control lines used in this study were previously reported as CC1 (now C2) and *CHD2* WT/WT2 (now C3) from (Tidball *et al.*, 2016; Tidball *et al.*, 2017), respectively, and were reprogrammed using identical methods.

##### CRISPR genome-editing with simultaneous reprogramming

P2 fibroblasts were reprogrammed as described above except with the addition of two "nickase" (D10A) CRISPR/Cas9 plasmids (px462) containing gRNAs flanking the P2 variant with a 39 bp offset. A single-stranded oligo donor nucleotide (ssODN) template was added with a silent point mutation engineered to disrupt the PAM sequence of one of the gRNAs (see **Supplemental Table 1** for gRNA, ssODN, and primer sequences). For the first 2 days of culture, the non-homologous end joining inhibitor, L755,507 was added to the medium at 2 μM (Yu *et al.*, 2015). Colonies were picked and genomic DNA sequenced as previously described (Tidball *et al.*, 2018). One line was generated with only the wild-type allele; however, the silent mutation did not incorporate. Subsequent long-range PCR determined the mutant allele had a ~500 bp deletion causing the apparent loss on the shorter PCR reaction (Primers found in **Supplemental Table 1** and DNA gel image in **Supplemental Figure 3**). This hemizygous line was used to determine whether loss of the mutant allele rescued phenotypes in the patient cells.

#### qPCR for integration of reprogramming vectors

Genomic DNA was isolated using the DNeasy kit (Qiagen). Using primers for the plasmid specific WPRE region, we performed qPCR on all iPSC lines to assay for integration of the reprogramming vectors. Quantitative PCR was performed using Power Sybr Green Master Mix (Thermo Fisher) as 20 μL reactions with 0.5 μM concentration of each primer and 10 ng of genomic DNA. Similar reactions for actin genomic DNA were performed to normalize to the genomic DNA content. Primer sequences are found in **Supplemental Table 1**.

#### SNP Chip Microarray and g-band karyotyping

Genomic DNA samples were isolated using the DNeasy kit (Qiagen). These samples were submitted to the University of Michigan Sequencing Core. A HumanCoreExome-24 v1.1 Infinium Whole genome genotyping BeadArray (Illumina) was performed on these samples and analyzed for copy number variations (CNVs) using KaryoStudio software (Illumina). iPSC lines were also submitted for standard g-band karyotype analysis to Cell Guidance Systems. For each line, 20 metaphase spreads were analyzed. Cell lines with karyotype abnormalities or significant CNVs (limit of detection average of ~ 0.15 MB) not also identified in the source fibroblasts were excluded from this study.

#### Excitatory neuron differentiation

We utilized a modification of the Shi *et al.* technique for differentiating iPSCs into excitatory cortical neurons (Shi *et al.*, 2012). iPSC lines were passaged using Accutase (Innovative cell) and replated onto Matrigel-coated 6-well plates at 2-3 x $10^5$ cells/mL in mTeSR1 with 10 μM rho-kinase inhibitor (Y-27632; Tocris, 1254). The medium without the inhibitor was replaced daily until the cells reach 80-100% confluency. The medium was then changed to 3N (50:50 DMEM/F12:neurobasal with N2 and B27 supplements) (Shi *et al.*, 2012) without vitamin A with 2 µM DMH1 (Tocris, 4126), 2 µM XAV939 (Cayman Chemical, 13596) , and 10 µM SB431542 (Cayman Chemical, 13031) (4 mL of medium per well). This medium was changed daily with 1 µM cyclopamine (Cayman Chemical, 11321) added beginning on day 1. On day 8, the cells were passaged using Dispase and gentle trituration to generate small clumps of neuroepithelium. These clumps were centrifuged at 50 g for 2 min to remove cell debris and single cells. The clumps were then resuspended in 3N medium with vitamin A and plated at 1:6 dilution onto fresh Matrigel-coated dishes. The 3N medium was changed daily for 14 days. Neural progenitors were then passaged using Accutase onto PEI/laminin coated dishes and coverslips at 2.5 x $10^5$ cells/mL. The day after passaging the medium was changed to BrainPhys with 20 ng/mL brain-derived neurotrophic factor (BDNF; Peprotech) and 20 ng/mL glial-derived neurotrophic factor (GDNF; Peprotech) with 0.2 µM Compound E (EMD Biosciences) and 0.4 mM dibutyral-cAMP (Sigma). Half volume medium changes were performed 3 times per week with compound E and dbcAMP only used during the first week after replating. The neuronal cultures were transduced with lentivirus containing a mouse *Camk2a* promoter-driven GFP to identify mature neurons (Shcheglovitov *et al.*, 2013; Nehme *et al.*, 2018). Cells were used for electrophysiological recordings between days 21-28 after final replating (43-50 days total differentiation).

#### Generation of stable doxycycline (dox)-inducible *Ngn1/2* iPSC lines

Established iPSC lines were incubated for 8 min in Accutase and 3x$10^5$ cells were replated in one well of a Matrigel-coated 6-well plate in mTeSR1 medium with 10 µM of the rho-kinase Y27632 inhibitor. After 24 h, the cells were transfected using Mirus LT-1 with 2 TALEN plasmids (pZT-C13-R1 and pZT-C13-L1, gifts of Jizhong Zou; addgene.org: #52638 and 52637, respectively) targeting the safe-harbor-like locus, *CLYBL,* and a targeting plasmid containing the dox-inducible promoter system which controls *Neurogenin1* (*Ngn1*) and *Ngn2* expression, and constitutively active *mCherry* and puromycin resistance genes (pUCM-CLYBL-*Ngn1/2*, a gift of Dr. Michael Ward). Medium was changed daily with fresh mTeSR1. After 5-7 days, clonal patches of *mCherry* positive cells were manually picked using an inverted epifluorescence microscope in a HEPA filtered workstation. These clones are typically heterogeneously mixed with non-integrated iPSCs. Positive selection during subsequent passages was used until the line was 100% mCherry-positive.

#### Differentiation of induced neurons (iNeurons)

Approximately $2x10^5$ cells/mL were plated into each well of a 6-well plate (Matrigel-coated) after accutase dissociation with Y27632 in mTeSR1 media. The mTeSR1 media was replaced daily with mTeSR1 media containing 1 ug/ml dox to generate iNeurons. After 72 hours of dox exposure, accutase was used to replate the cells as a single cell suspension at $2x10^5$ cells/mL onto PEI/laminin coated dishes in 3N medium containing 1 μg/mL dox. Medium was changed daily with fresh 3N medium containing dox. On day 8, the medium was replaced with BrainPhys medium containing N2 and SM1 supplements as well as 20 ng/ml BDNF and 20 ng/ml GDNF. Half-media changes were performed 3 times/week.

#### Electrophysiological recordings

iPSC-derived neurons were analyzed by whole-cell patch clamp recordings. Voltage clamp recordings were performed in the standard whole-cell configuration, using previously described conditions (Liu *et al.*, 2013). Isolated $I_{Na}$ was recorded from single GFP+ cells (labeled with a lentiviral *Camk2a*-GFP reporter) at RT (21–22°C) in the presence of a bath solution containing (in mM): 120 NaCl, 1 $BaCl_2$, 2 $MgCl_2$, 0.2 $CdCl_2$, 1 $CaCl_2$, 10 HEPES, 20 TEA-Cl, and 10 glucose (pH = 7.35 with CsOH; osmolarity = 300–305 mOsm). Fire-polished patch pipettes were filled with an internal solution containing (in mM): 1 NaCl, 150 N-methyl-D-glucamine, 10 ethylene glycol tetra acetic acid (EGTA), 2 $MgCl_2$, 40 HEPES, and 25 phosphocreatine-tris, 2 MgATP, 0.02 $Na_2GTP$, 0.1 leupeptin (pH = 7.2 with $H_2SO_4$). All recordings were performed within 10 to 120 min after the culture medium was replaced by bath recording solution and the dish with cells was placed on the recording setup. Cells with peak $I_{Na}$ density < 60 pA/pF were excluded from analysis due to presumed lack of maturity or cell integrity.

For current-clamp recordings of APs in iPSC-derived neurons, patch pipettes were filled with internal solution consisting of (in mM): 140 K-gluconate, 4 NaCl, 0.5 $CaCl_2$, 10 HEPES, 5 EGTA, 2 Mg-ATP, and 0.4 GTP (pH = 7.2, adjusted with KOH). iPSC-derived neurons were bathed in BrainPhys media during current clamp recordings. Single APs were evoked from the resting membrane potential by injection of a series of depolarizing pulses 1 ms long with 0.01 nA increments beginning at subthreshold levels and rising until APs were consistently generated. The threshold potential for initiation of AP, depolarization rate, peak amplitude, amplitude half-width, and repolarization rate were measured using the Event Detection tool in Clampfit program. The threshold potential was defined as the level with membrane potential change of >10 mV/ms when the first AP was initiated. The rise rate was measured as changes in membrane potentials during the time from threshold potential to the AP peak. Because the repolarization phases of APs from patient iPSC-neurons were mostly very slow and took a long time to reach the resting membrane potential level, we arbitrarily measured the repolarization potentials at 5 ms, 10 ms and 40 ms post-peak. Repetitive AP firing was evoked by injection of a series of 1500 ms currents varying from -60 pA to 180 pA in 10 pA steps from resting membrane potential. Cells with resting membrane potentials more depolarized than -47 mV were excluded due to presumed lack of maturity or cell integrity. For repetitive evoked spiking analysis, cells with a maximum firing frequency of less than 4 AP/s were excluded. Signals were amplified with a Multiclamp 700B amplifier (Molecular Devices, Sunnyvale, CA) and filtered at 2-4 kHz and digitized at 20 kHz for offline analysis. Access resistances were usually less than 10 MΩ with 70% series resistance compensation and monitored throughout the experiments to ensure their constancy. Data were acquired with a Digidata 1440A interface and analyzed using pClamp10 offline. All experiments were carried out at RT (21-22 °C).

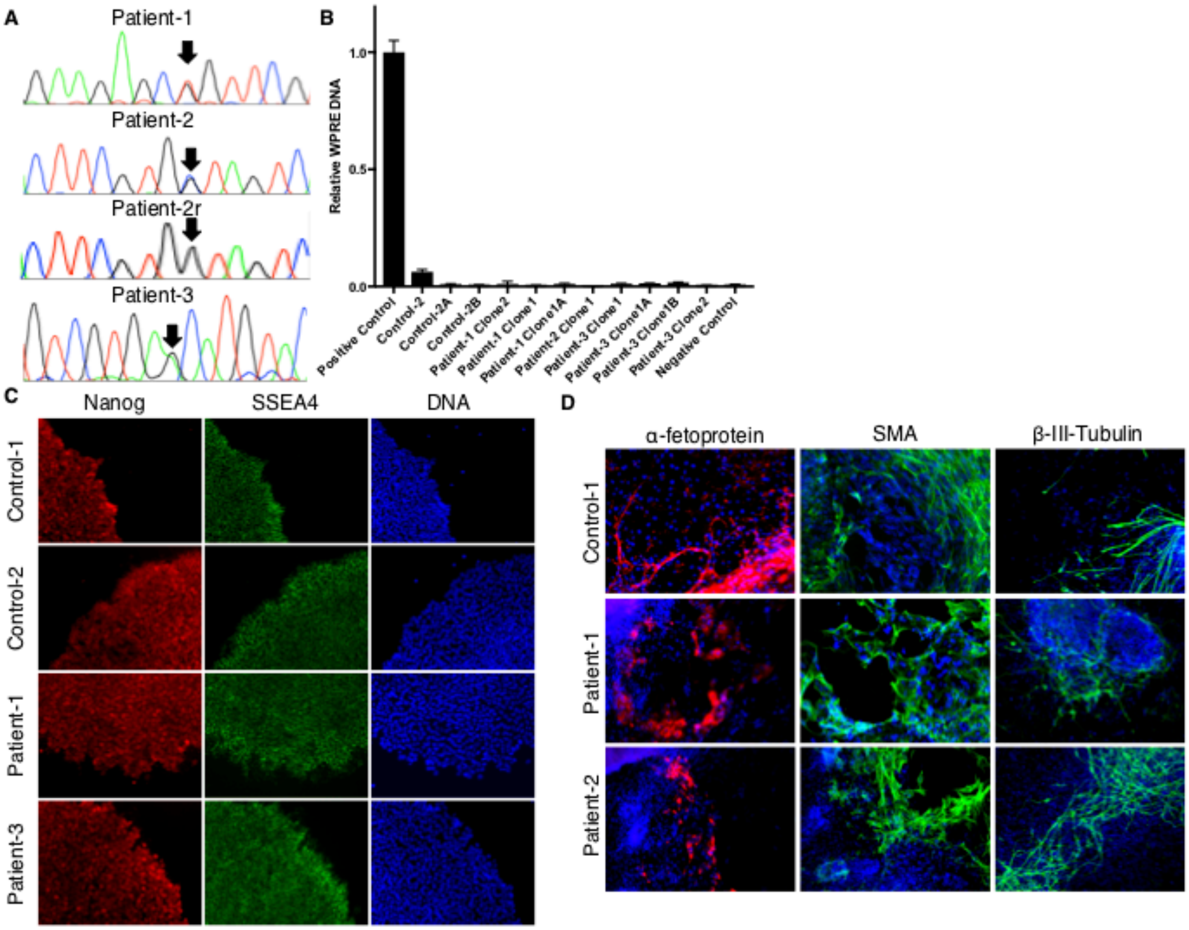


**Supplemental Figure 1. Validation of control and patient iPSC lines.** (**A**) PCR products from each of the three patients over the mutated region from fibroblast lines were Sanger sequenced. The chromatographs showing the heterozygous missense point mutations for each patient are highlighted with an arrow. (**B**) Quantitative PCR for the WPRE elements was performed for all iPSC lines used as compared to a positive control with validated integration of the episomal plasmids. (**C**) Immunostaining for pluripotency markers NANOG and SSEA4, which are not on the reprogramming plasmids. (**D**) Embryoid bodies treated with FBS were plated and stained for markers of the three germ layers α-fetoprotein (endoderm), smooth muscle actin (mesoderm), and β-III-tubulin (neuroectoderm).

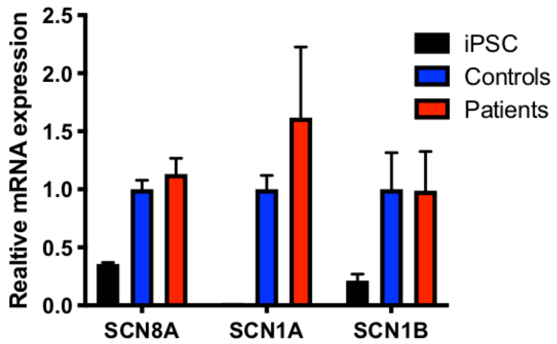


**Supplemental Figure 2. No differences in VGSC transcript levels between controls and patients**. Total RNA was isolated from patient and control iPSC-derived neurons that had been differentiated and plated for 21-28 days. cDNA generated using oligoDT primers was used as template for qPCR for VGSC genes (*SCN8A, SCN1A,* and *SCN1B*). ΔΔCT was performed using actin as the endogenous control. iPSC N = 2. Control N = 6 (C3 = 3, C4 = 3). Patient N = 8 (P1 = 1, P2 = 2, and P3 = 5).

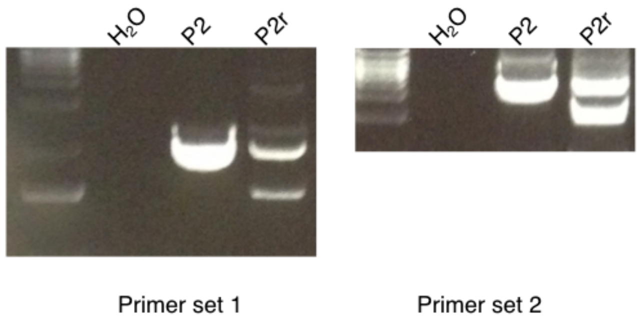


**Supplemental Figure 3. Patient-2 rescue line contains large deletion of disease allele.**

Two PCR primer sets were used for long-range PCR around the CRISPR cut site (2067 and 4642 bp products). In both cases, a ~500bp smaller PCR product was observable in the P2 CRISPR line indicating a large deletion rather than the presumed disease allele correction.

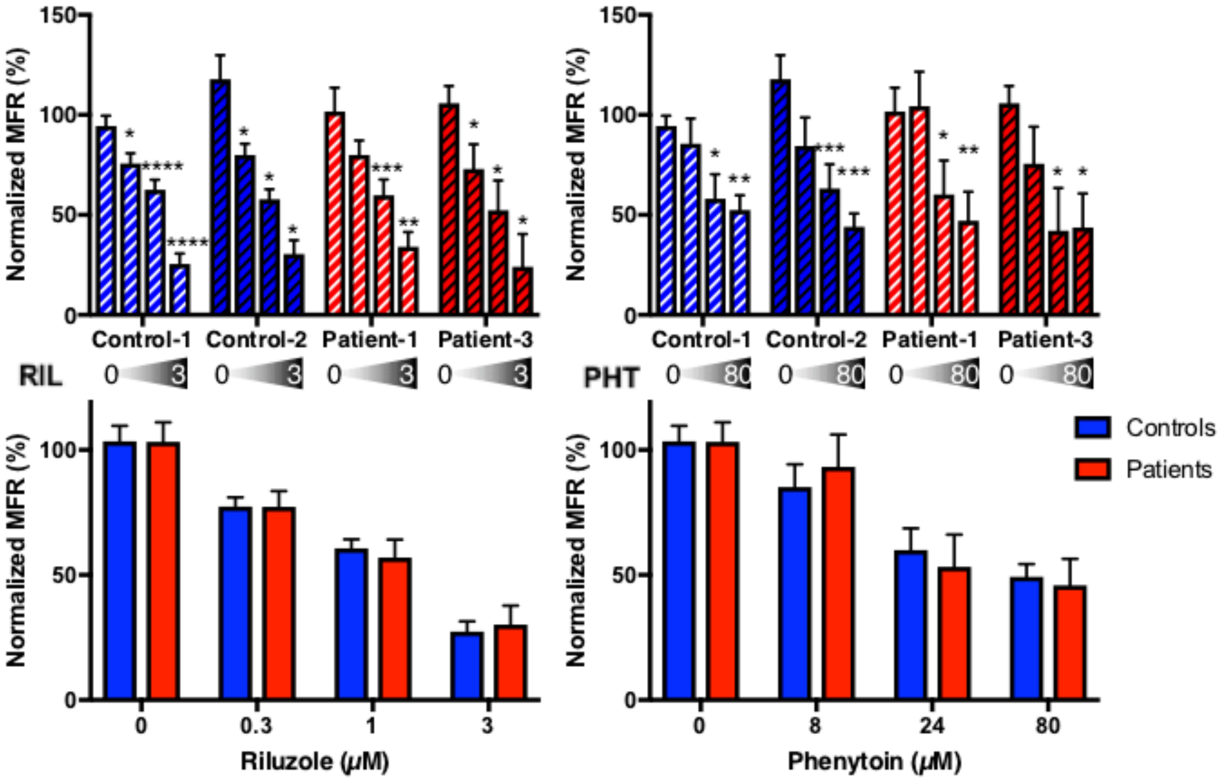


**Supplemental Figure 4. Riluzole and phenytoin reduce MFR but not in a genotype dependent manner.** (**A, B**) Individual patient neuron normalized mean firing rate response to increasing concentrations of riluzole (0, 0.3, 1, and 3 μM) and phenytoin (0, 8, 24, and 80 μM), respectively. N = 8 independent experiments for C2 and P1. N = 5 independent experiments for C3 and P3. Within group comparisons for drug dosage effect compared to vehicle (DMSO) were performed using a one-way ANOVA. (**C, D**) Combined effects of controls and patients for riluzole and phenytoin, respectively. N = 13 for each group. Two-way unpaired ANOVA (due to missing values) was performed. All post-hoc analyses used the two-stage linear step-up procedure post-test. * < 0.05, ** < 0.01, *** < 0.001, and **** < 0.0001.

| Table S1. Primer and gRNA sequences. | |
|---|---|
| | **CRISPR Guide RNA Sequences** |
| SCN8A Patient-2 gRNA-1 | GCTAGTCATCCTCTCCATTG |
| SCN8A Patient-2 gRNA-2 | GTCGAAGATGTTCCAGCCAA |
| ssODN | TCTTCACCTGTGAGTGTGTGCTCAAAATGTTTGCGTTGAGGCACTACTACTTCACAATTGGCTGGAACATCTTCGACTTCGTGGTAGTCATCCTCTCCATTGTAGgtgagtggggttggggaagtaggCGAGGGGAAAGGGGACCTGACTGATTGTGAG |
| | **PCR Primers** |
| 1872 F | CAGACTTTGCAGATGCCTTG |
| 1872 R | GGCTCTTTCCCTTCTTCCTT |
| 1592 F | TCAGCAAGCCTTTGACATTG |
| 1592 R | GGGTATCTTGCCTGCACTTC |
| 1759 F | CACCCAGGGAGTGGCTTTAA |
| 1759 R | GCTCCAAGGCATCTGCAAAG |
| Actin qPCR F | CTGTGGCATCCACGAAACTA |
| Actin-qPCR R | AGCACTGTGTTGGCGTACAG |
| WPRE qPCR F | CTTCCCGTATGGCTTTCATT |
| WPRE qPCR R | GGGCCACAACTCCTCATAAA |
| *SCN8A* qPCR F | GAGATAGCGGGGAGTTGGAC |
| *SCN8A* qPCR R | GTGTGGTTGTGATTGGCTCG |
| *SCN1A* qPCR F | ATGGCCATGGAGCACTATCC |
| *SCN1A* qPCR R | CTACCAGGCTAAGCGTCACA |
| *SCN1B* qPCR F | GCCGTGTATGGGATGACCTT |
| *SCN1B* qPCR R | AGCGCAGGATCTTGACAAAC |
| Patient 2 long range 1 F | CCAGATTCCTGGCTTGAGAG |
| Patient 2 long range 1 R | AATCAACAGGGTGGCATTTC |
| Patient 2 long range 2 F | GTAGCTGGGCGTGGTTATGT |
| Patient 2 long range 2 R | GAGCAGTGGACAAGGGTAGC |

#### Supplemental Table 1.

A list of sequences for PCR primers, gRNAs, and ssODN.

| Table S2. Antibodies and dilutions | | | | |
|---|---|---|---|---|
| **Antigen** | **Species** | **Dilution** | **Vendor** | **Catalog#** |
| alpha-fetoprotein | Rabbit | 1:100 | Dako | A0008 |
| Ankyrin-G | Rabbit | 1:500 | Santa Cruz | sc-28561 |
| beta-III-Tub | Mouse | 1:1000 | Covance | MMS-435p |
| GABA | Rabbit | 1:5000 | Sigma | A2052 |
| MAP2ab | Mouse | 1:500 | Sigma | M2320 |
| Nanog | Rabbit | 1:500 | Abcam | ab21624 |
| SMA | Mouse | 1:1000 | Abcam | ab5694 |
| SSEA4 | Mouse | 1:200 | DSHB | MC-813-70 |
| vGlut1 | Guinea Pig | 1:200 | Millipore | AB5905 |
| anti-Rabbit AlexaFluor594 | Goat | 1:500 | Invitrogen | A-11037 |
| anti-Mouse AlexaFluor488 | Goat | 1:500 | Invitrogen | A-11029 |
| anti-Guinea Pig AlexaFluor647 | Goat | 1:500 | Invitrogen | A-21450 |

**Supplemental Table 2.**

A list of antibodies and dilutions used in this study.

| Table S3. Passive properties | | | | |
|---|---|---|---|---|
| Cell Line | Genotype | Resting Vm, mV | Input Resistance, MΩ | Capacitance, pF |
| Control-1 (1 clone) | WT | 53.5 ± 1.1 (8) | 1500 ± 300 (7) | 8.6 ± 1.0 (7) |
| Control-2 (1 clone) | WT | 53.3 ± 1.7 (6) | 680 ± 210 (6) | 9.0 ± 1.6 (3) |
| Patient-1 (1 clone) | R1872>L | 50.4 ± 0.9 (7) | 910 ± 230 (7) | 7.1 ± 0.6 (7) |
| Patient-2 (2 clones) | V1592>L | 50.7 ± 2.3 (3) | 2000 ± 200 (3) | 5.2 ± 2.2 (2) |
| Patient-3 (1 clone) | N1759>S | 50.6 ± 0.9 (10) | 620 ± 90 (9) | 7.5 ± 1.6 (8) |
| Control | WT | 53.4 ± 0.1 (2) | 1100 ± 60 (2) | 8.8 ± 0.3 (2) |
| Patient | Mutant | 50.6 ± 0.2 (3)* | 1200 ± 700 (3) | 6.6 ± 1.2 (3) |
| Control-2B (1 clone) | WT | 55.6 ± 0.8 (36) | 407 ± 35 (36) | 9.9 ± 0.8 (33) |
| Control-3C (1 clone) | WT | 57.5 ± 1.2 (20) | 449 ± 47 (20) | 10.7 ± 1.0 (20) |
| Patient-1A (1 clone) | R1872>L | 55.7 ± 0.5 (59) | 313 ± 25 (59) *** | 10.3 ± 0.7 (59) |
| Patient-3A (1 clone) | N1759>S | 54.4 ± 0.9 (35) | 580 ± 66 (34) | 9.0 ± 0.5 (32) |
| Control iNeuron | WT | 56.5 ± 0.9 (2) | 453 ± 45 (2) | 10.3 ± 0.4 (2) |
| Patient iNeuron | Mutant | 55.1 ± 0.7 (2) | 449 ± 134 (2) | 9.6 ± 0.7 (2) |

**Supplemental Table 3.**

Passive neuronal properties of resting membrane potential, input resistance, and capacitance are quantified for all neurons used in this study.

| Table S4. Voltage dependent properties | | | | | | |
|---|---|---|---|---|---|---|
| | **Activation Curve** | | | **Inactivation Curve** | | |
| | **$G_{max}$ (nS)** | ***k (mV)*** | **$V_{1/2}$ (mV)** | ***h* (mV)** | **$V_{1/2}$ (mV)** | **C** |
| Control | 3.32±0.70 | 5.00±0.45 | -35.3±1.8 | -4.98±0.25 | -53.6±1.6 | 0.02±0.004 |
| Patient 1 | 1.99±0.23 | 5.80±.023 | -34.9±2.0 | -4.84±0.26 | -52.7±1.2 | 0.03±0.006 |
| Patient 2 | 3.07±0.91 | 4.70±0.32 | -34.4±1.9 | -5.14±0.49 | -57.0±2.4 | 0.03±0.007 |
| Patient 3 | 1.62±0.37 | 5.36±0.47 | -32.2±1.4 | -4.12±0.31 | -50.9±1.2 | 0.03±0.006 |

**Supplemental Table 4.**

Voltage-dependent properties for control and patient dual-SMAD neuron activation and inactivation curves.

### References

Liu Y, Lopez-Santiago LF, Yuan Y, Jones JM, Zhang H, O'malley HA*, et al.* Dravet syndrome patient-derived neurons suggest a novel epilepsy mechanism. Annals of neurology 2013; 74(1): 128-39.

Nehme R, Zuccaro E, Ghosh SD, Li C, Sherwood JL, Pietilainen O*, et al.* Combining NGN2 Programming with Developmental Patterning Generates Human Excitatory Neurons with NMDAR-Mediated Synaptic Transmission. Cell reports 2018; 23(8): 2509-23.

Okita K, Matsumura Y, Sato Y, Okada A, Morizane A, Okamoto S*, et al.* A more efficient method to generate integration-free human iPS cells. Nature methods 2011; 8(5): 409-12.

Shcheglovitov A, Shcheglovitova O, Yazawa M, Portmann T, Shu R, Sebastiano V*, et al.* SHANK3 and IGF1 restore synaptic deficits in neurons from 22q13 deletion syndrome patients. Nature 2013; 503(7475): 267.

Shi Y, Kirwan P, Smith J, Robinson HP, Livesey FJ. Human cerebral cortex development from pluripotent stem cells to functional excitatory synapses. Nature Neuroscience 2012; 15(3): 477-86.

Tidball AM, Dang LT, Glenn TW, Kilbane EG, Klarr DJ, Margolis JL*, et al.* Rapid generation of human genetic loss-of-function iPSC lines by simultaneous reprogramming and gene editing. Stem cell reports 2017; 9(3): 725-31.

Tidball AM, Neely MD, Chamberlin R, Aboud AA, Kumar KK, Han B*, et al.* Genomic instability associated with p53 knockdown in the generation of huntington's disease human induced pluripotent stem cells. PloS one 2016; 11(3): e0150372.

Tidball AM, Swaminathan P, Dang LT, Parent J. Generating loss-of-function iPSC lines with combined CRISPR indel formation and reprogramming from human fibroblasts. Bio Protoc 2018; 8.

Yu C, Liu Y, Ma T, Liu K, Xu S, Zhang Y*, et al.* Small molecules enhance CRISPR genome editing in pluripotent stem cells. Cell stem cell 2015; 16(2): 142-7.
